# Stochastic inference of clonal dominance in gene therapy studies

**DOI:** 10.1101/2022.05.31.494100

**Authors:** L. Del Core, M. A. Grzegorczyk, E. C. Wit

## Abstract

Clonal dominance is a wake-up-call for adverse events in gene therapy applications. This phenomenon has mainly been observed as a consequence of a malignancy progression, and, in some rare cases, also during normal haematopoiesis. We propose here a random-effects stochastic model that allows for a quick detection of clonal expansions that possibly occur during a gene therapy treatment.

Starting from the Ito-type equation, the dynamics of cells duplication, death and differentiation at clonal level without clonal dominance can be described by a local linear approximation. The parameters of the base model, which are inferred using a maximum likelihood approach, are assumed to be shared across the clones. In order to incorporate the possibility of clonal dominance, we extend the base model by introducing random effects for the clonal parameters. This extended model is estimated using a tailor-made expectation maximization algorithm. The main idea of this paper is to compare the base and the extended models in high dimensional clonal tracking datasets by means of Akaike Information Criterion in order to detect the presence of clonal dominance. The method is evaluated using a simulation study, and is applied to investigating the dynamics of clonal expansion in a in-vivo model of rhesus macaque hematopoiesis.

**Author summary:** Preventing or quickly detecting clonal dominance is an important aspect in gene therapy applications. Over the past decades, clonal tracking has proven to be a cutting-edge analysis capable to unveil population dynamics and hierarchical relationships in vivo. For this reason, clonal tracking studies are required for safety and long-term efficacy assessment in preclinical and clinical studies. In this work we propose a random-effects stochastic framework that allows to investigate events of clonal dominance using high-dimensional clonal tracking data. Our framework is based on the combination between stochastic reaction networks and mixed-effects generalized linear models. We have shown in a simulation study and in a real world application that our method is able to detect the presence of clonal expansions. Our tool can provide statistical support to biologists in gene therapy surveillance analyses.

## Introduction

The idea of gene therapy is that the correction of the defective gene(s) underlying the disease is, in principle, sufficient for inducing disease remission or even full recovery [1]. Since the blood system possesses a hierarchical structure with haematopoietic stem cells (HSC) at its root [2], correction of HSCs might be sufficient to eradicate a genetic disease in the blood system. Since transduction of stem cells has proven to be less efficient than transduction of more mature cells, it might be necessary to allow very high gene transfer rates, i.e., multiple vector copies, to ensure efficient genetic modification of HSCs [3, 4]. But genetic modification of large numbers of cells is associated with the higher probability of unintentional vector insertion events near the growth-regulatory genes that may lead to insertional mutagenesis [5–7]. One particular drawback of insertional mutagenesis is the phenomenon of clonal dominance, which occurs if one or more clones dominate cell production [8]. The most extreme case of clonal dominance is monoclonality, where an entire tissue is dominated by the progeny of one particular cell. Although different gradients of clonal dominance (oligoclonality) exist, a precise threshold that defines dominance is hard to be specified in general, and thus a clear definition of what is meant by clonal dominance is required for any particular study.

Clonal dominance in malignant haematopoiesis has been previously identified as a consequence of a clonal competition that is corrupted by disease progression [9, 10]. However, clonal dominance has also been observed in normal haematopoiesis, even in the case of truly neutral clonal markers [11–13]. Indeed, on the basis of various mathematical models, progression of monoclonality has been discussed also for normal (non-leukaemic) stem cell systems [14–18]. While there is strong evidence for clonal selection inducing monoclonal systems in the crypts of the small intestine [19–22], such a process has not been demonstrated for the haematopoietic system yet. To shed more light on those mechanisms, in this manuscript we extend the work of [23, 24] and propose a random-effects cell differentiation network to model the dynamics of clonal expansions for high dimensional clonal tracking data.

More in detail, starting from the definition of the master equation [25], a set of Ito stochastic differential equations is derived to describe the first-two-order moments of the process. We estimate the parameters of the Ito system from its Euler-Maruyama local linear approximation (LLA) [26]. We propose a new inference procedure in the LLA formulation using a maximum likelihood approach, replacing the iterative weighted least square algorithm previously developed in [23, 24]. Although the base LLA model formulation has been shown to be effective in modelling cell differentiation [24], it has some limitations as it only provides an average description of the dynamics across all the clones, and does not take into account possible extreme behaviour. Indeed in the base model all the dynamics parameters are shared across the clones, and thus is not possible to identify heterogeneous clonal patterns. Thus the base LLA formulation cannot be used to model clonal dominance. Therefore in this work we further increase the flexibility of the base LLA model to check if the process dynamics is mainly due to few clones and if those dominate a particular cell type. To this end we introduce random effects for the clones inside the LLA formulation, providing a mixed-effects LLA model. Then, if the mixed model outperforms the fixed one in terms of Akaike Information Criterion we use the former to infer the process parameters in order to identify which clones are mainly expanding and in which cell compartments. As every mixed-effects formulation, inference of parameters is performed by means of an expectation-maximization algorithm, for which we developed an efficient implementation. Effectively, our random-effects LLA formulation describes a stochastic process of clonal dominance on a network of cell lineages. We tested and validated our method using a simulation study. Finally, our model allowed to investigate the dynamics of clonal expansion in a in-vivo model of rhesus macaque hematopoiesis [27].

## Materials and Methods

This section contains background on clonal tracking data and a description of a Rhesus Macaques study. We also provide a concise description of the clonal dominance model and inference procedure. A more comprehensive, mathematical description can be found in Section 1 of S1 Text.

### Clonal tracking data

There are several high-throughput systems capable to quantitatively track cell types repopulation from an individual stem cell after a gene therapy treatment [28–30]. Tracking cells by random labeling is one of the most sensitive systems [31]. In gene therapy applications, haematopoietic stem cells (HSCs) are sorted from the bone marrow of the patient and uniquely labeled by the random insertion of a viral vector inside its genome. Each label, called clone, vector integration site (VIS), or barcode, is defined as the genomic coordinates where the viral vector integrates. After transplantation, all the progeny deriving through cell differentiation inherits the original labels. During follow-up, the labels are collected from tissues and blood samples using Next Generation Sequencing (NGS) [32–35]. Therefore NGS does allow identifying, quantifying and tracking clones arising from the same HSC ancestor. Over the past decades, clonal tracking has proven to be a cutting-edge analysis capable to unveil population dynamics and hierarchical relationships in vivo [36–39].

We consider single cell barcode data collected from an established hematopoietic stem cell gene therapy model previously used to investigate the hematopoietic reconstitution in Rhesus Macaques. [27] applied a lentiviral cellular barcoding technology to rhesus CD34+ HSPCs, thus allowing clonal tracking after myeloablative autologous transplantation. In particular, mobilized peripheral blood (MPB) CD34+ cells from three macaques were transduced with barcoded vectors, and 7.8-16.7 million autologous GFP+ cells were reinfused after an ablative total body irradiation. Following engraftment, myeloid Granulocytes (G), Monocytes (M), and lymphoid T, B, and Natural Killer (NK) cells were flow sorted (purity median 98.8%).

The authors showed with high confidence (> 95%) that a single barcode marked only one HSPC clone at these transplanted doses [40, 41]. Thus, only a minority of clones containing more than one barcode would skew calculations of the frequency of repopulating clones upward, but would not impact analysis of lineage contributions or kinetics. Barcode retrieval by PCR, Illumina sequencing, and custom data analysis was performed on purified hematopoietic lineage samples monthly for 9.5 months (ZH33), 6.5 months (ZH17), and 4.5 months (ZG66) [42]. They demonstrated high reproducibility of barcode retrieval and quantitation via sequencing several replicates on the same collected DNA samples. They also assayed independently processed replicate blood samples to identify a lower barcode read threshold that would result in 95% barcode retrieval between replicates. In particular they established a sampling error threshold of 1144 reads. Therefore we also considered the same reads threshold here, so as to be consistent with the previous studies. The total numbers of clones collected are 1165 (`ZH33`), 1280 (`ZH17`), and 1291(`ZG66`). To further remove bias, we only focused on the clones recaptured at least 5 times across lineages and time. This resulted in a subset of clones of size 481(`ZH33`), 139 (`ZH17`), and 202 (`ZG66`). Further details on transduction protocols and culture conditions can be found in the original study.

### A stochastic model for cell differentiation

We consider three event types, such as cell duplication, cell death and cell differentiation for a time counting process

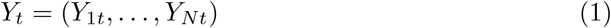

of a single clone in *N* distinct cell lineages. The time counting process *Y*_*t*_ for a single clone in a time interval (*t, t* + Δ*t*) evolves according to a set of reactions {*v*_*k*_}_*k*_ and hazard functions {*h*_*k*_}_*k*_ defined as

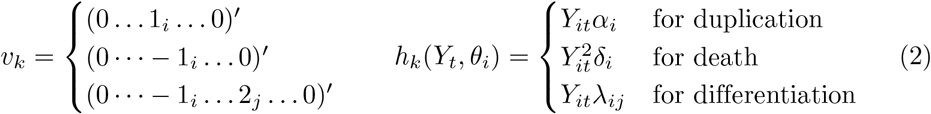

which contains a linear growth term with a duplication rate parameter *α*_*i*_ > 0, a quadratic term for cell death with a death rate parameter *δ*_*i*_ > 0, and a linear term to describe cell differentiation from lineage *i* to lineage *j* with differentiation rate λ_*ij*_ > 0 for each *i* ≠ *j* = 1,…, *N*. Finally we use the LLA formulation of Section 1.3.1 from S1 Text with net-effect matrix and hazard vector defined as

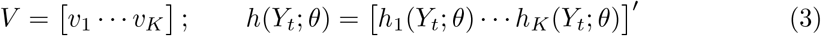

In this formulation we implicitly assume that cells belonging to the same lineage obey to the same dynamics laws, that is all the clones share the same vector parameter *θ*. In case we argue that clones behave differently in terms of dynamics we can use the random-effects LLA formulation of Eq. (6), where the random effects are defined on the vector parameter *θ* w.r.t. the clones. This is the case in our application study presented in next section, where we check wether there is heterogeneity in the clones for the duplication and death parameters, which we use as a proxy for a clonal expansion or contraction.

### LLA formulation of clonal dominance

Let Y_*t*_ = (*Y*_1*t*_,…, *Y*_*Nt*_) be a collection of “cells” of *N* different types at time *t* obeying to a network of stochastic biochemical reactions defined by a net-effect matrix *V* ∈ ℤ^*N*×*K*^, a vector parameter *θ* and an hazard vector *h*(*Y, θ*) = (*h*_1_(*Y, θ*),…, *h*_*K*_(*Y, θ*)) and let

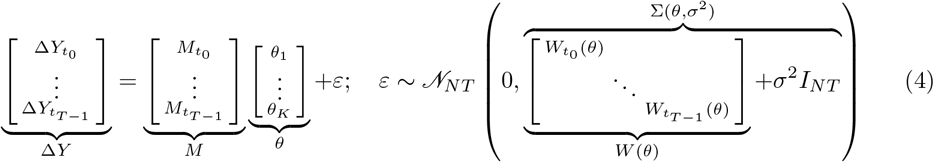

be the local linear approximation of an Ito-type equation written in generalized linear model formulation (see Section 1 of S1 Text for details) where

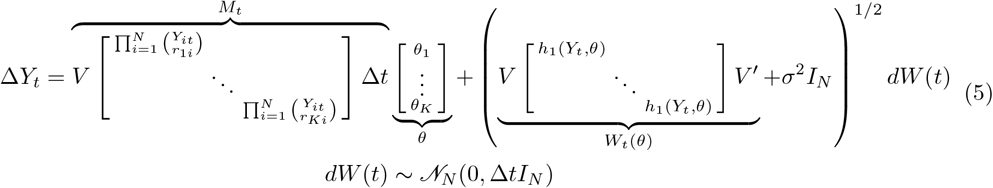

*σ*^2^ is the noise variance, *M*_t_*θ* the mean drift, *W*_*t*_(*θ*) the diffusion matrix, and Δ*Y*_*t*_ = *Y*_*t*+Δ*t*_ − *Y*_*t*_ is a finite-time increment. In system (4) all the cell counts *Y*_1_,…, *Y*_*N*_ share the same parameter vector *θ*. To infer the parameters of (4)-(5) we developed a maximum likelihood algorithm which is fully described in Section 1.4 of S1 Text. In some cases it may happen that the cells being analysed are drawn from a hierarchy of *J* different populations that possibly behave differently in terms of dynamics. In this case it might be of interest to quantify the population-average *θ* and the subject-specific effects *u* around the average *θ* for the description of the subject-specific dynamics. Therefore we introduce here a novel stochastic framework which is more flexible then the base LLA model thus allowing for the quantification of clonal contribution to the process. In particular, to quantify the contribution of each subject *j* = 1,…, *J* on the process dynamics we extend the LLA (4) with a mixed-effects model [43] introducing random effects *u* for the *J* distinct subjects on the parameter vector *θ*, leading to a random-effects stochastic reaction network (`RestoreNet`). The extended random-effects formulation becomes

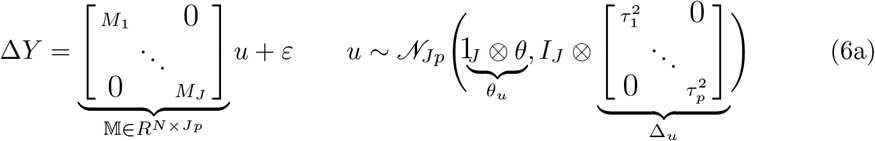

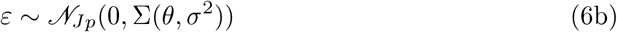

where 𝕄 is the block-diagonal design matrix for the random effects *u* centered in *θ*, where each block *M*_*j*_ is subject-specific. As in the case of the null model (4), to explain additional noise of the data, which has the additional advantage of avoiding singularity of the covariance matrix *W* (*θ*), we add to its diagonal a small quantity *σ*^2^ which we infer from the data. Under this framework it can be shown that

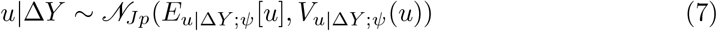

where 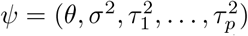 is the set of all the unknown parameters. Once the parameters are estimated (see next section for inference details), the conditional expectations *E*_*u*|Δ*Y*;ψ_[*u*] can then be used as a proxy for the clone-specific rate parameters. This method allows to infer the clone-specific dynamic by extremely reducing the problem dimensionality from *J* · *p* to 2 · *p* + 1 (*J* ≫ 2).

### Inference procedure

In order to infer the Maximum Likelihood estimator 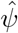 for 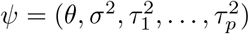 we develop an efficient tailor-made Expectation-Maximization algorithm where the collected cell increments Δ*Y* and the random effects *u* take the roles of the observed and latent states respectively. The full analytical expression of *E*_*u*|Δ*Y*;*ψ*_[*u*], *V*_*u*|Δ*Y* ;*ψ*_ (*u*), the `E`-step function *Q*(*ψ*|*ψ*^*^) = *E*_*u*|*ΔY* ; *ψ*_* [𝓁(Δ*Y, u*;*ψ*)] and its partial derivatives 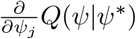 are available (see Section 1.4 of S1 Text). In the EM-algorithm we iteratively update the E-function *Q*(*ψ*|*ψ*^*^) using the current estimate *ψ*^*^ of *ψ* and then we minimize the −*Q*(*ψ*|*ψ*^*^) w.r.t. *ψ*. As the `E`-step function *Q*(*ψ*|*ψ*^*^) is non-linear, we used the `L-BFGS-B` algorithm from the optim() base `R` function for optimization, to which we provided the objective function, along with its gradient ∇_*ψ*_*Q*(*ψ*|*ψ*^*^), as input. Given the high-dimensionality of the clones being analysed, and due to the sparsity of the clonal tracking datasets, the `E`-step function *Q*(*ψ*|*ψ*^*^) and its gradient ∇_*ψ*_*Q*(*ψ*|*ψ*^*^) are written in a sparse block-diagonal matrix fashion, so as to reduce computational complexity and memory usage. The EM algorithm is run until a convergence criterion is met, that is when the relative errors of the E-step function *Q*(*ψ*|*ψ*^*^) and the parameters ^*ψ*\*^ are lower than a predefined tolerance.

Once we get the EM estimate 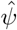 for the parameters we evaluate the goodness-of-fit of the mixed-model according to the conditional Akaike Information Criterion [44]. As every EM algorithm, the choice of the starting point *ψ*_*s*_ is very important from a computational point of view. We chose 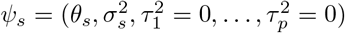 as a starting point where 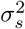 is the optimum found in the fixed-effects LLA formulation (4). This is a reasonable choice since we want to quantify how the dynamics 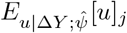 of each subject (clone) *j* departs from the average dynamics *θ*_*s*_. With the help of simulation studies (see Section 2 of S1 Text) we empirically proved that this choice always led to a conditional expectation *E*_*u*|Δ*Y* ;*ψ*_[*u*] consistent with the true clone-specific dynamic parameters *θ*. Computational details can be found in Section 1.4 of S1 Text. The pseudocode of the EM algorithm is provided in Algorithm 3 of S1 Text.

### Computational implementation

The maximimul likelihood inference for the basal model and the expectation maximization algorithm for the random-effects model are implemented in the 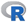 package `RestoreNet`. Few minimal working examples showing the usage of the package are provided in Section 5 of S1 Text.

## Results

A first comparative evaluation study on synthetic data, whose results are provided in Section 2 of S1 Text, shows how the proposed random-effects formulation is able to identify clonal dominance. We found that the random-effects model reached a significantly lower AIC than the null model, thus detecting the simulated dominance of a single clone into a cell type.

Next, we compared the base and random-effects models on the clonal tracking data of the rhesus macaque study fully described in Section “Clonal tracking data”. Although the sample DNA amount was maintained constant during the whole experiment (200 ng for `ZH33` and `ZG66` or 500 ng for `ZH17`), the sample collected resulted in different magnitudes of total number of reads (see Table 2 from S1 Text). This discrepancy makes all the samples not comparable across time and cell types. Therefore we rescaled the barcode counts according to Eq. (19) from S1 Text, and we report the rescaled cell counts, at clonal level, in Fig. 1. Since the CD34+ cells were not collected, we only estimated the duplication parameters *α*_*T*_, *α*_*B*_, *α*_*NK*_, *α*_*M*_, *α*_*G*_ and the death parameters *δ*_*T*_, *δ*_*B*_, *δ*_*NK*_, *δ*_*M*_, *δ*_*G*_ of the lymphoid (T, B, NK) and myeloid (M, G) cells. Therefore the differentiation parameters are not present in our model, and the net-effect matrix and the hazard vector are obtained from Eq. (2) - (3) accordingly. The corresponding model becomes effectively a birth/death model. We fitted both the fixed model (4) and the mixed-effects model (6) separately to the data of each animal, where *J* is equal to 481 (`ZH33`), 139 (`ZH17`), and 202 (`ZG66`) respectively. The size of the dynamic vector parameter *θ* is equal to 10, that is one scalar value for each combination of the five cell types with the duplication and death reactions. Also, *N* equals 11275 (`ZH33`), 2555 (`ZH17`), and 2770 (`ZG66`), while the number of time-points *T* is equal to 6 (`ZH33`), 5 (`ZH17`), and 4 (`ZG66`).

**Fig 1.**
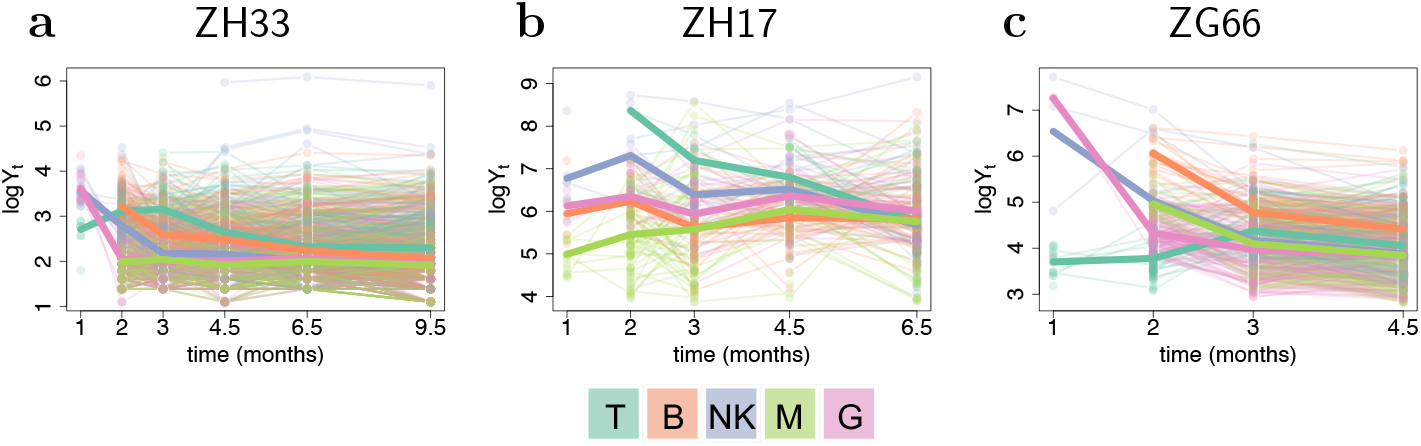
Logarithmic rescaled clonal abundance (*y*-axis) over time (*x*-axis) in each lineage (colors) of each treated animal (a-c). The thin lines are clone-specific, while the thick lines are the average across the clones. Data is rescaled according to equation (19) from S1 Text.

We report the results on model selection in Table 1 and the estimated parameters 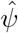 in Table 3 of Section 4 from S1 Text. Then, from the estimated parameters 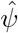 following Eq. (18) from S1 Text we computed the conditional expectations 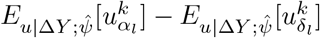, which we use as a proxy for the *k*-th clone-specific net-duplication *α*_*l*_ − *δ*_*l*_ in each cell lineage *l*. The resulting values are reported in Fig. 2 in a box-plot fashion. To visualize our findings at clonal level, in Fig. 3 we propose to use a weighted pie chart. Each pie corresponds to a particular clone and is weighted by the corresponding conditional expectations 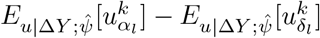. The biological interpretation of this figure is that the larger the diameter, the more the corresponding clone is dominating cell production into the lineage associated to the largest slice.

**Table 1.**
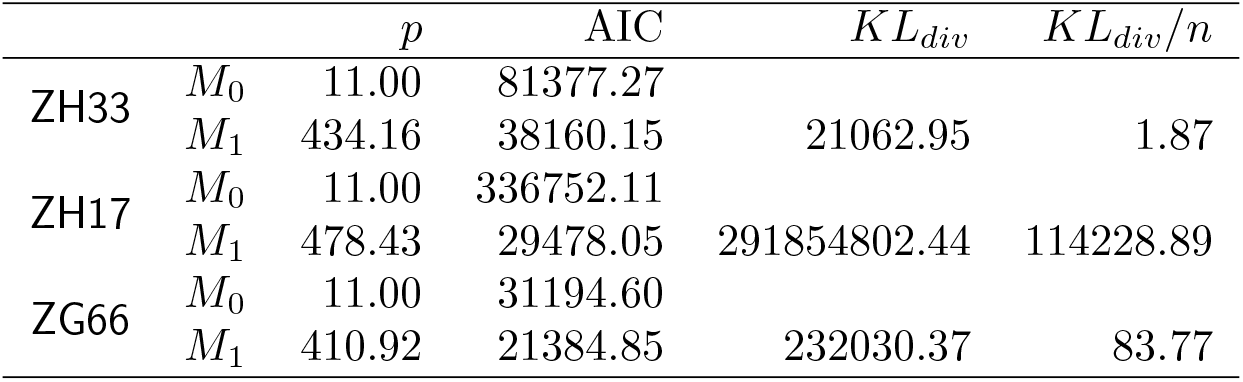
Comparison between fixed and mixed effects model: Number of parameters (*p*), Akaike Information Criterion (AIC), Kullback-Leibler divergence (*KL*_*div*_) and rescaled *KL*_*div*_ (*KL*_*div*_/*n*) for the fixed (*M*_0_) and the mixed (*M*_1_) models in each treated individual.

| | | $p$ | AIC | $KL_{div}$ | $KL_{div}/n$ |
| --- | --- | --- | --- | --- | --- |
| ZH33 | $M_0$ | 11.00 | 81377.27 | | |
| | $M_1$ | 434.16 | 38160.15 | 21062.95 | 1.87 |
| ZH17 | $M_0$ | 11.00 | 336752.11 | | |
| | $M_1$ | 478.43 | 29478.05 | 291854802.44 | 114228.89 |
| ZG66 | $M_0$ | 11.00 | 31194.60 | | |
| | $M_1$ | 410.92 | 21384.85 | 232030.37 | 83.77 |

**Fig 2.**
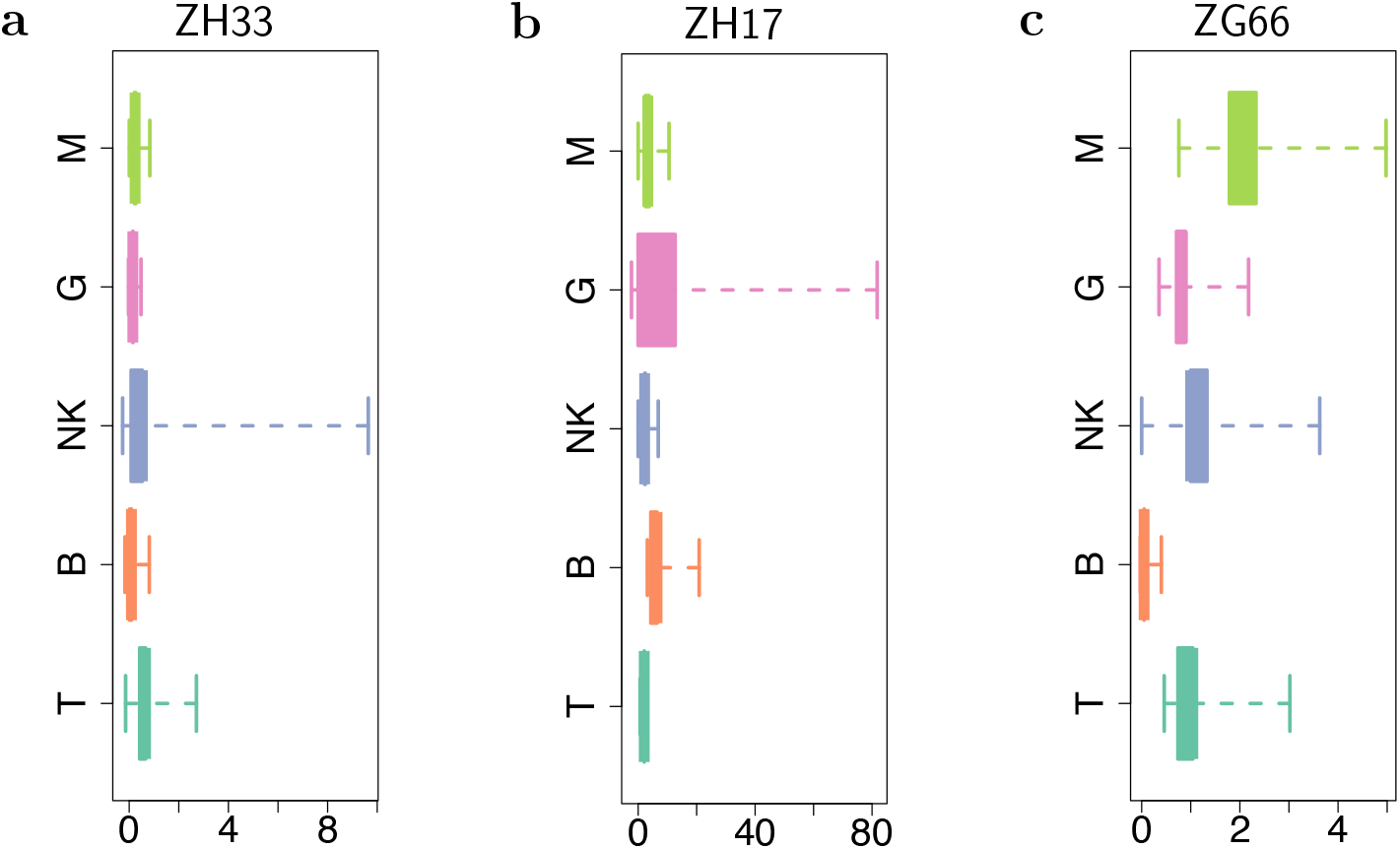
For each animal analyzed (a-c), the boxplots of the conditional expectations 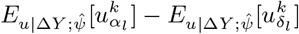 computed from the estimated parameters 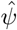 for the clone-specific net-duplication *α*_*l*_ − *δ*_*l*_ in each cell lineage *l* (different colors). The whiskers extend to the data extremes.

**Fig 3.**
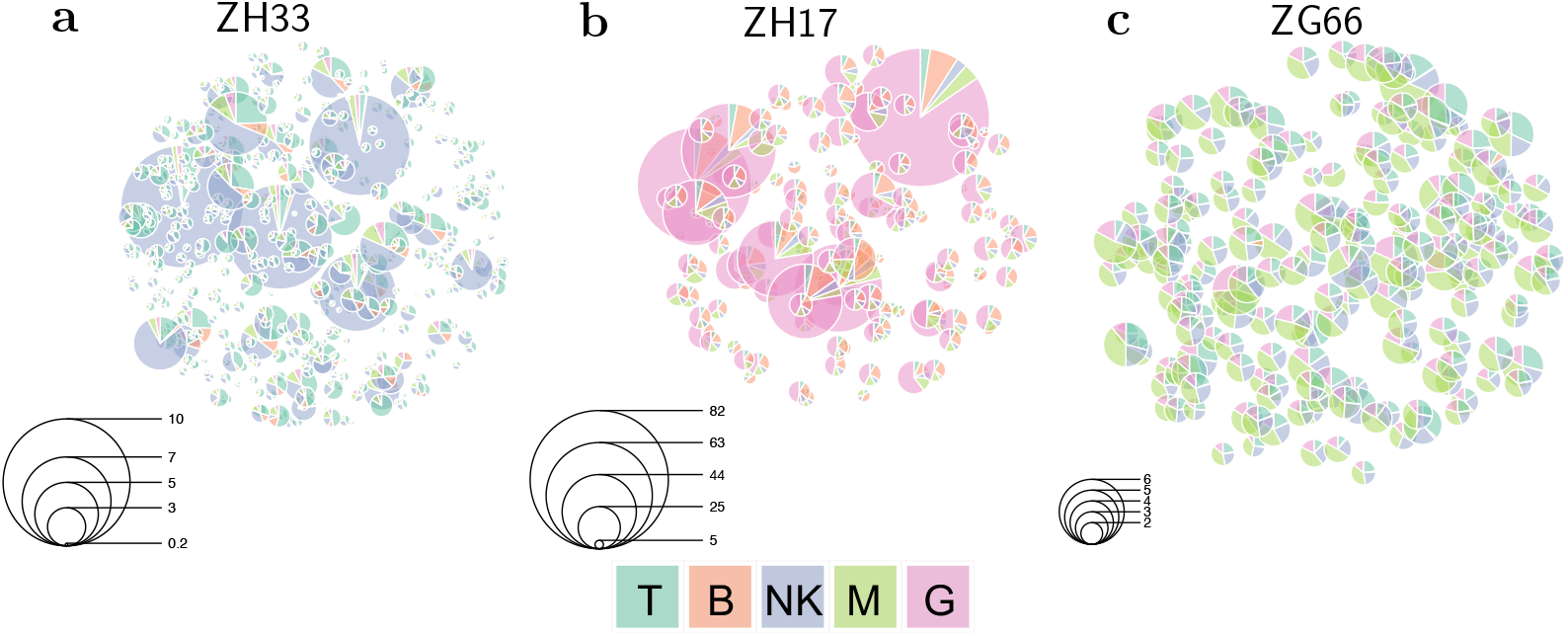
Graphical representation of the results obtained with the proposed the mixed effects model: Random effects for the clones to the process parameters of the rhesus macaques ZH33 (a), ZH17 (b) and ZG66 (c). Each *k*-th clone is identified with a pie whose slices are lineage-specific and weighted with *w*_*k*_ defined as the difference between the corresponding duplication and death parameters, that is 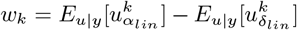. The diameter of the *k*-th pie is proportional to the euclidean 2-norm of *w*_*k*_. The legend scales are different across the three plot panels.

As a result, according to the AIC values, in each treated individual the mixed model (*M*_1_) outperformed the fixed one (*M*_0_). This means that the clones did not follow the same average dynamics for the birth/death process. Instead, the dynamic of some clones departed from the average dynamics with a significant (random) effect. In particular, the conditional net-duplication rates 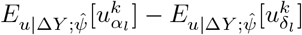 of Fig. 2 -3 suggest that there is clonal dominance in specific cell lineages. As an example, for the animals `ZH33` and `ZG66` we observed clonal expansions into `NK` cells with high conditional rates. Whereas, for the animal `ZH17` we observed clonal expansions into `G` and `B` cell lineages with high conditional rates. Finally, for the animal `ZG66` we also observed events of clonal dominance into `M` and `T` cell lineages. Furthermore, the weighted pie charts shown in Fig. 3 reveal different gradients of clonal dominance between the three rhesus macaques. As an example, looking at the size of the pies it is possible to observe an higher clonal dominance of `NK` cells in `ZH33` and of `G` cells in `ZH17` compared to the expansions of `M`, `NK` and `T` cells detected in `ZG66`, where the diameters of the clone-specific pies are rather similar. Not only does the proposed mixed effect model detect clonal dominance of certain cell types, it is also able to detect which clones are responsible.

## Discussion and conclusion

In this work we proposed a random-effects cell differentiation network which takes into account heterogeneity in the dynamics across the clones. Our framework extends the clone neutral local linear approximation of a stochastic quasi-reaction network, written in the Ito formulation, by introducing random-effects for the clones to allow for clonal dominance. To infer the parameter of the base (fixed-effects only) model we used a maximum likelihood approach. Whereas, to infer the parameters of the random-effects model, we have developed an expectation-maximization (EM) algorithm. We tested our framework with a *τ*-leaping simulation study (see Section 2 from S1 Text for details), showing accurate performance of the method in the identification of a clonal expansion and in the inference of the true parameters. Subsequently, the application of the method on a rhesus macaque clonal tracking study revealed significant clonal dominance for specific cell types. Particularly interesting is that the `NK` clonal expansions detected by our model were already observed by former studies [27,45,46], and therefore our findings are consistent with those previously obtained. Indeed [45] described the oligoclonal expansions of NK cells and the long-term persistence of HSPCs and immature `NK` cells.

The main approximation in both the basal and random-effects formulations is the piece-wise linearity of the process. That is, in both cases we consider first a local linear approximation of the Ito equation, which then we use to infer the process parameters either with or without random-effects. Although the linearity assumption makes all the computations easier, this approximation becomes poor as the time lag increments (the δ*ts*) of the collected data increase. This can be addressed by introducing in the likelihood higher-order approximation terms than the ones considered by the Euler-Maruyama method. The Milstein approximation is a possible choice. Another, completely different, approach is to employ extended Kalman filtering (EKF) which is suitable for non-linear state space formulations. Also, our framework cannot consider false-negative errors or missing values of clonal tracking data. Also for this issue, an EKF formulation could be a possible extension.

Our tool can be considered as complementary to the classical Shannon entropy index [47] in detecting fast and uncontrolled growing of clones after a gene therapy treatment. Indeed, while the Shannon entropy measures the diversity of a population of clones as a whole, `RestoreNet` provides a clone-specific quantification of dominance in terms of conditional mean and variance of the expansion rates. In conclusion, our proposed stochastic framework allows to detect deviant clonal behaviour relative to the average dynamics of hematopoiesis. This is an important aspect for gene therapy applications where is crucial to quickly detect clonal dominance to prevent any adverse event that may be related to malignant scenarios. Therefore our tool can provide statistical support to biologists in gene therapy surveillance analyeses. With slight modifications our framework can be applied to every study of population dynamics that can be described with an Ito-type formulation, even when the whole population needs to be drawn from an hierarchical structure having subject-specific dynamics.

### S1 Text Stochastic inference of clonal dominance in gene therapy studies

(PDF)

## Author Contributions

- All authors analysed the data and wrote the paper.
- L.D.C. designed and implemented the stochastic framework.

## Data availability

The data that supports the findings of this study is openly available at [27].

## Code availability

- The code that supports the findings of this study is openly available at https://github.com/delcore-luca/ClonalDominance
- The stochastic framework is implemented in the 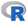 package `RestoreNet` available at https://github.com/delcore-luca/RestoreNet

## Acknowledgements

- This publication is based on work from COST Action CA15109 (COSTNET), supported by COST (European Cooperation in Science and Technology).
- E.C.W. acknowledges support from the Fondazione Leonardo (514.7.010.098-4) and funding from the Swiss National Science Foundation (SNSF 188534)
- The funders had no role in study design, data collection and analysis, decision to publish, or preparation of the manuscript.

## S1 Text: Stochastic inference of clonal dominance in gene therapy studies

### 1 Mathematical details

#### 1.1 Stochastic quasi-reaction networks

Stochastic quasi-reaction networks (S-QRN) allow to implement a particular class of stochastic differential equations that can be used to model biochemical reactions. More formally, let

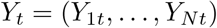

be a collection of molecules of *N* different types observed at time *t*, and consider *K* distinct (and competing) reactions

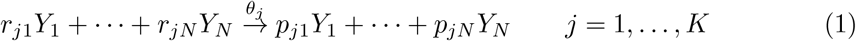

each occurring with its own rate *θ*_*j*_. The coefficients *r*_*ji*_’s defining the left-side of the reaction are called reagents and represent the minimum amount of molecules of type *i* needed for the *j*-th reaction to occur. Similarly, the coefficients *p*_*ji*_ defining the right-side of the reaction are called products and represents the amount of produced molecules of type *i* after the *j*-th reaction is triggered. We assume that, if we observe *Y*_0_ = (*r*_*j*1_,…, *r*_*jN*_) molecules at time *t* = 0, the *j*-th reaction will occur after

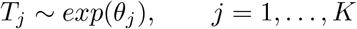

Namely, if exactly *r*_*ij*_ molecules of each type *i* would be present, then the *j*-th reaction can only take place in one way, with the exponential hazard rate *θ*_*j*_. The interpretation is that, after a waiting time *T*_*j*_, *r*_*ji*_ molecules of type *i* collide with each other and produce *p*_*ji*_ molecules of type *i* (∀*i* = 1,…, *N*), while the molecules move randomly in a hosting “cellular” environment. However, in general at time *t* = 0 we might observe *Y*_*i*0_ ≥ *r*_*ji*_ molecules of each type *i* and, therefore, the *j*-th reaction can take place in a combinatorial number of ways leading to the following waiting time formulation

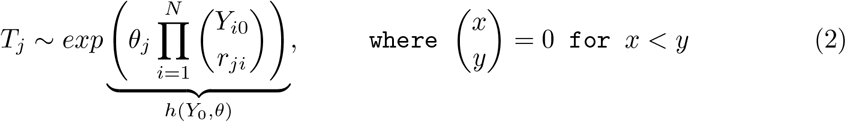

In this case, the effect will be that at time *t* + *T*_*j*_ we have the following expression for the number of molecules of substrate *i*,

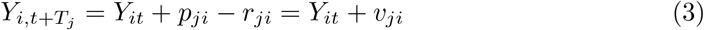

where *v*_*ji*_ = *p*_*ji*_ − *r*_*ji*_ is the *j*-th net effect. More compactly, for a set of *K* reactions and *N* species, the molecular transfer from reagent to product species is a net change of

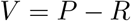

##### Algorithm 1: *τ* -leaping algorithm

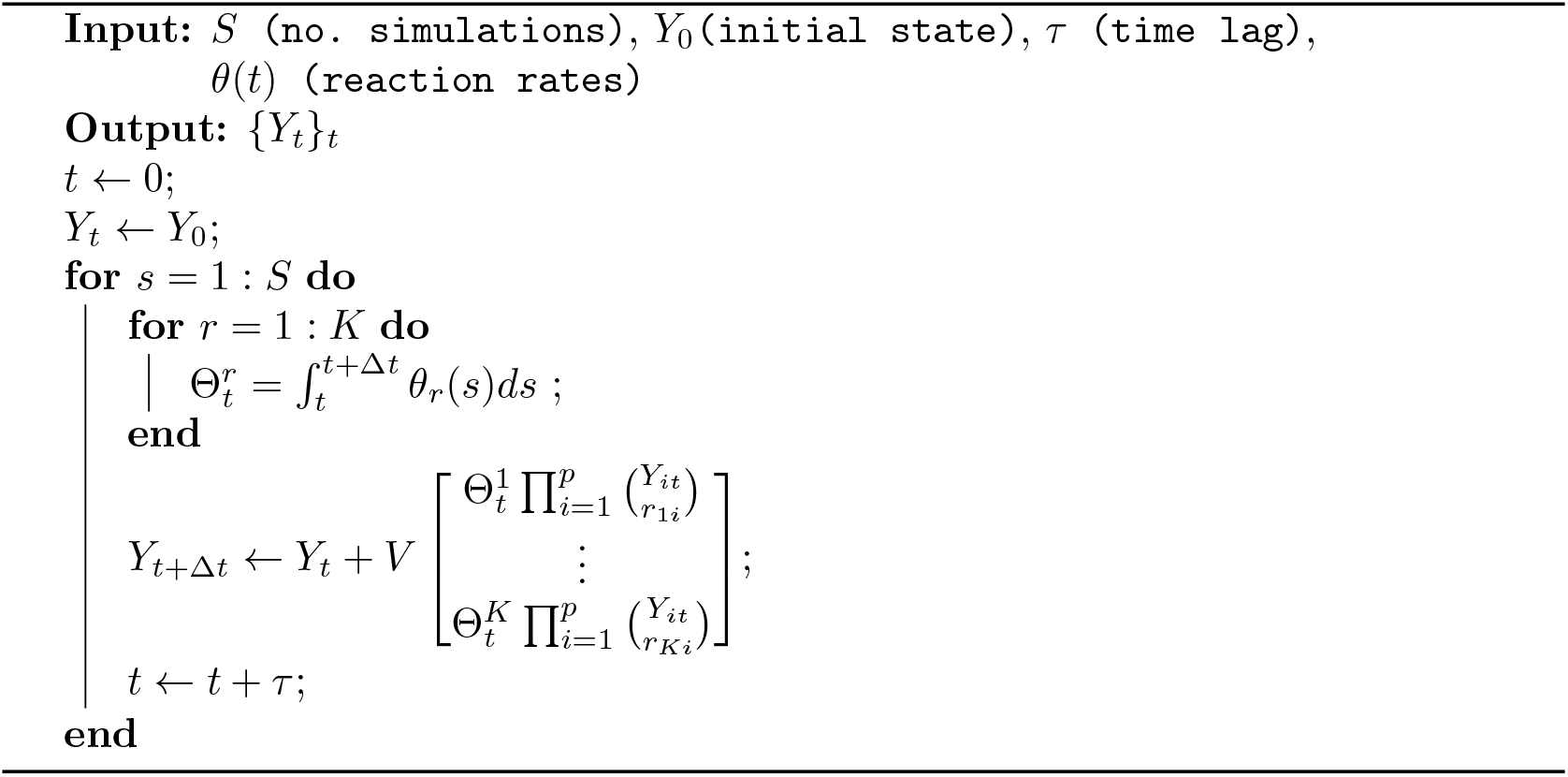

where *P* = [*p*_*ji*_]′ denotes the *N* × *r* dimensional matrix of products, *R* = [*r*_*ji*_]^′^ is the *N* × *r* dimensional matrix of reactants, and *V* = [*v*_*ji*_]^′^ is an *N* ×*r* dimensional matrix called net-effect matrix. Therefore, a S-QRN of *K*-distinct reactions is fully identified by a net-effect matrix *V* and by the hazard vector

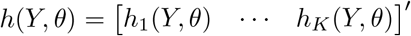

#### 1.2 Simulating a trajectory of molecules

A *τ*-leaping algorithm is an alternative method to a Gillespie algorithm for simulating triggering-chain events. Instead of simulating a waiting time for the first reaction to occur and selecting the corresponding winner reaction, a *τ*-leaping algorithm simulates the number of occurrences of each possible event after a time-lag equal to *τ* elapsed. Formally, let { *N*_*r*_(*t*)}_*t*≥0_ be an inhomogeneous Poisson point process representing the number of reactions of type *r* that took place up to (and including) time *t*. Therefore

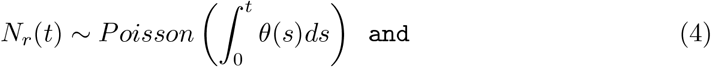

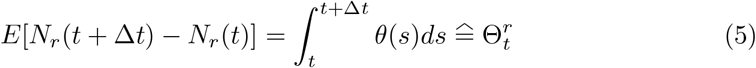

The last equation gives an estimate of the expected number of reactions of type *r* that took place within the time interval [*t, t* +. Δ_t_[. Furthermore, the expected number of molecules *Y*_*t*+Δ*t*_ at time *t* + Δ*t* given the current number of molecules *Y*_*t*_ is given by

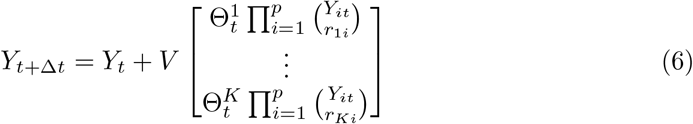

The pseudocode of the *τ*-leaping algorithm (*τ*-`LA`) is reported in Algorithm 1.

#### 1.3 Inference of the rates

##### 1.3.1 Local linear approximation

In order to estimate the rates *θ* = [*θ*_1_ … *θ*_*K*_]′, we focus on the first two-order moments of the process, that is we consider the Ito equation

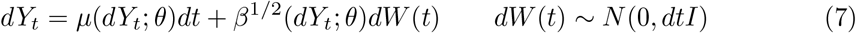

where *dY*_*t*_ = *Y*_*t*+*dt*_ - *Y*_*t*_ is an infinitesimal time drift and *μ*(*dY*_*t*_; *θ*) and *β* (*dY*_*t*_; *θ*) are called mean-drift and diffusion respectively. Given a *σ*-algebra (Ω, *ℱ*, ℙ), the solution

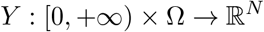

of (7) is called Ito diffusion. Instead of finding the Ito diffusion itself, we focus on the first two-order moments *μ*(*dY*_*t*_; *θ*) and *β* (*dY*_*t*_; *θ*) of the infinitesimal time drift *dY*_*t*_ which can be approximated with the following Lemma and Proposition.

###### Lemma 1.

*Given the hazard function as a limit of a conditional probability*

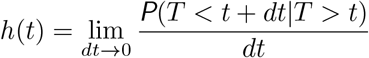

*for small dt the following approximation holds*

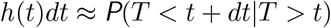

*Furthermore, the event* {*Y*_*t+dt*_ − *Y*_*t*_ ≡ *V*_·*j*_} ≡ {*Y*_*t+dt*_ = *Y*_*t*_ + *V*_·*j*_} *occurs with probability P*(*T*_*j*_ <*t* + *dt*|*T*_*j*_ > *t*).

###### Proposition 1.

*An approximation of the mean drift μ*(*dY*_*t*_; *θ*) *and the diffusion β* (*dY*_*t*_; *θ*) *for a small time increment dt is given by*

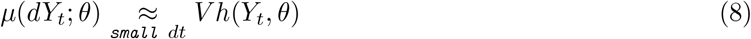

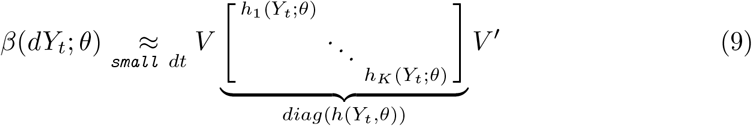

*Proof*.

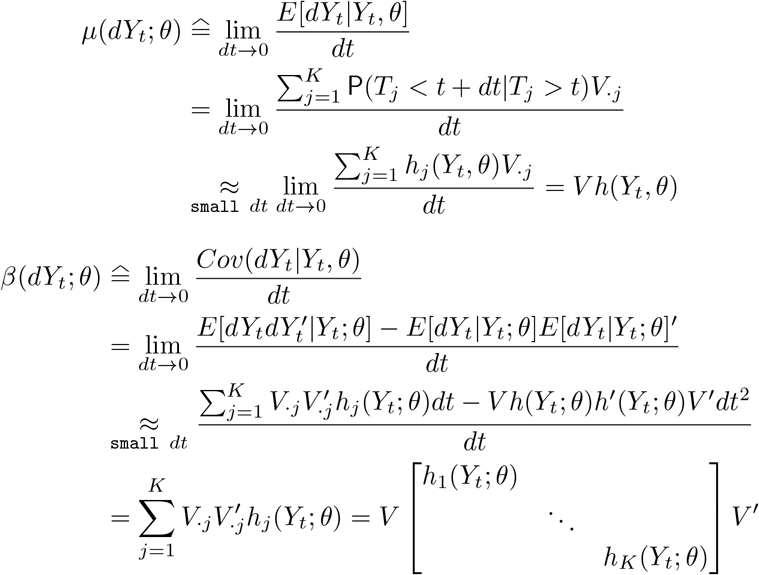

Using previous results and some linear algebra, the approximated Ito equation (7) can be further approximated as

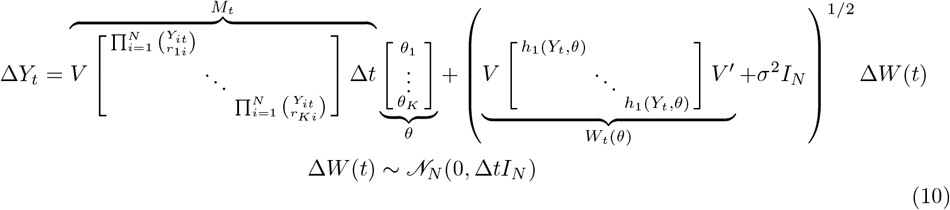

or more compactly

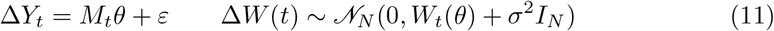

where we included the term *σ*^2^*I*_*N*_ so as to prevent singularity of the diffusion term, and to additionally explain noise variance. In practice, since we collect only discrete-time increments. Δ*Y*_*t*_ = *Y*_*t*+Δ*t*_ − *Y*_*t*_, we consider an Euler-Maruyama local linear approximation (LLA) of the approximated Ito equation. Indeed we also replaced the infinitesimal increments *dt* and *dY*_*t*_ with the discrete increments. Δ*t* and. Δ*Y*_*t*_. Then, all the time-specific blocks can be stacked together obtaining the full generalized linear model (GLM) formulation

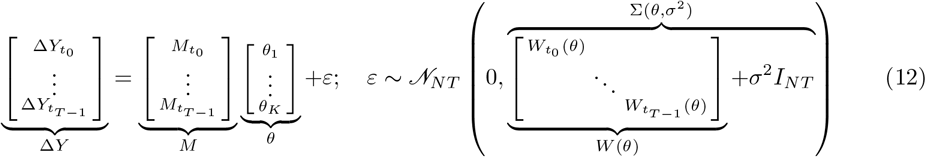

which is convenient for parameters inference.

##### 1.3.2 Maximum Likelihood (ML)

We infer the parameters (*θ, σ*^2^) with a maximum likelihood approach, that is we solve the following constrained optimization problem

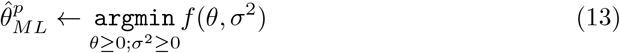

where the objective function is

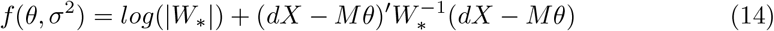

and we compactly write the diffusion matrix *W*_*_ = *W*(*θ, σ*^2^) as a function of the free parameters. Using the rules of matrix calculus [1], the partial derivatives of *f* w.r.t. *θ* and *σ*^2^ can be written as

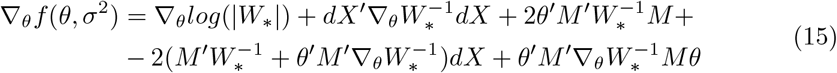

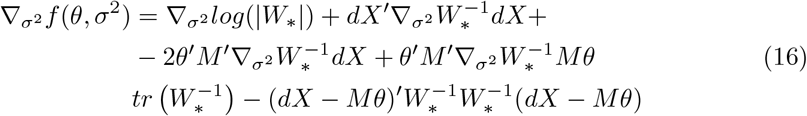

where

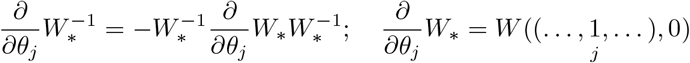

#### Algorithm 2: Maximum Likelihood inference for the base (null) model

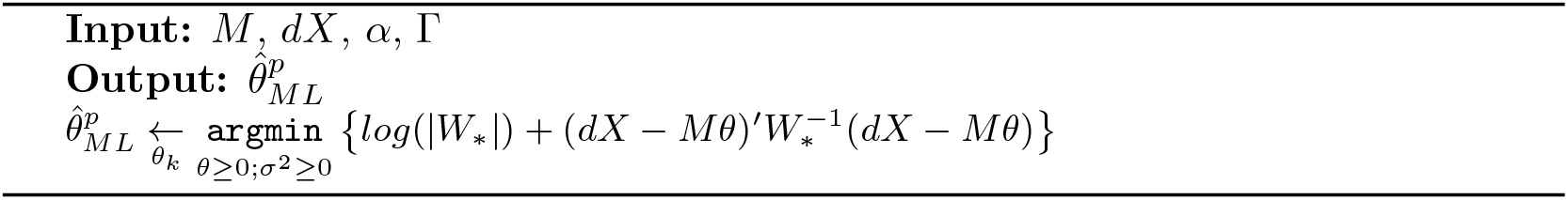

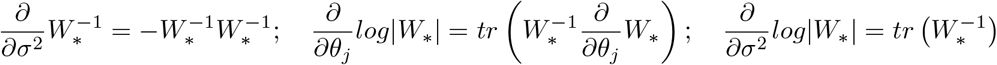

Then, we solve (26) by using the objective function (14) and its gradients (15)-(16) inside the `L-BFGS-B` optimization algorithm from the `optim()` function of the `stats` `R` package. The inference procedure is summarised in Algorithm 2.

#### 1.4 A mixed effects LLA model

In the system (24) all the molecules *Y*_*1*_,…, *Y*_*N*_ share the same parameter vector *θ*. In some cases it may happen that the molecules being analysed are drawn from a hierarchy of *J* different populations having different properties. In this case it might be of interest to quantify the population-average *θ* and the subject-specific effects *u* around the average *θ* for the description of the subject-specific dynamics. Therefore, to quantify the contribution of each subject *j* = 1,…, *J* on the process’s dynamics we extended the LLA (24) by introducing random effects *u* for the *J* distinct subjects on the parameter vector *θ*, leading to the following mixed-effects [2] formulation

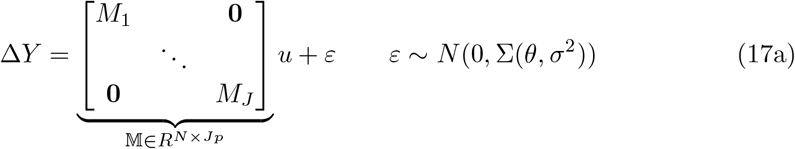

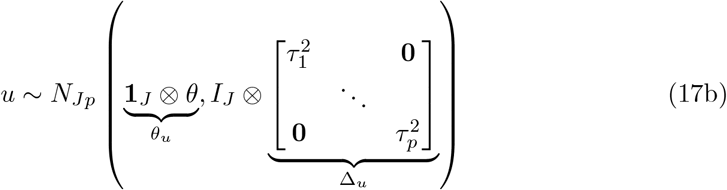

where 𝕄 is the block-diagonal design matrix for the random effects **u** centered in *θ*, where each block *M*_*j*_ is subject-specific. As in the case of the null model (24), to explain additional noise of the data and to avoid singularity of the stochastic covariance matrix *W* (*θ*) we added to its diagonal a small unknown quantity *σ*^2^ which we infer from the data. In order to infer the maximum likelihood estimator 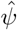 for 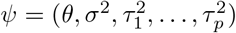 we developed an efficient tailor-made expectation-maximization algorithm where Δ*Y* and *u* take the roles of the observed and latent states respectively. Under this framework

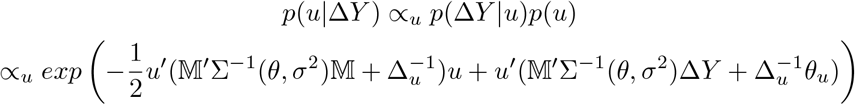

and therefore

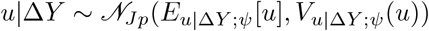

where

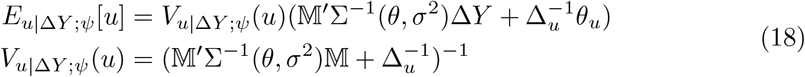

Also, the joint log-likelihood of Δ*Y* and *u* is given by

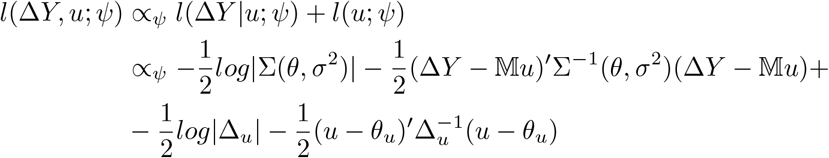

which only depends on *u* linearly via its first two-order conditional moments (18). Therefore, it follows for the `E`-step function that

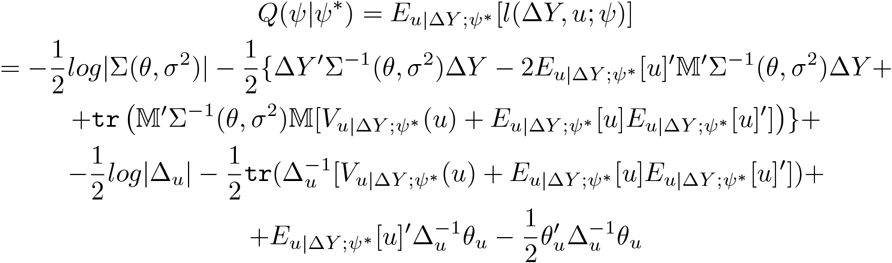

The gradient of Q(*ψ*| *ψ* ^*^) is defined by the following partial derivatives

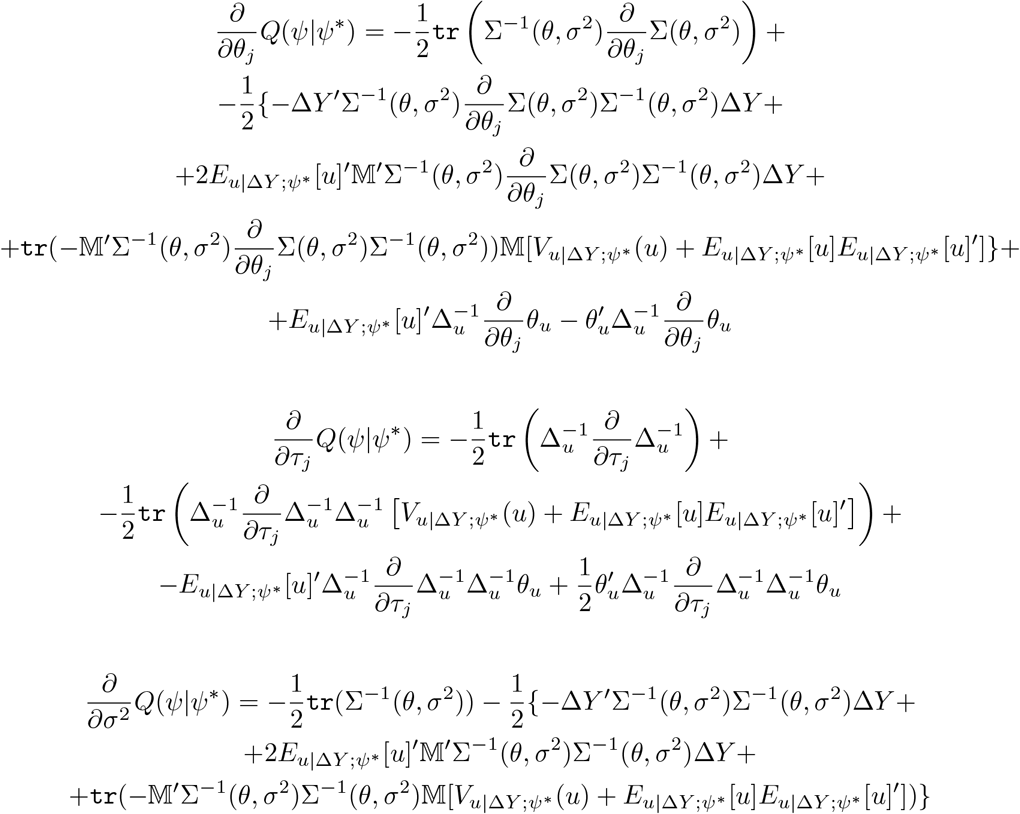

##### Algorithm 3: EM algorithm for the mixed-effects LLA model

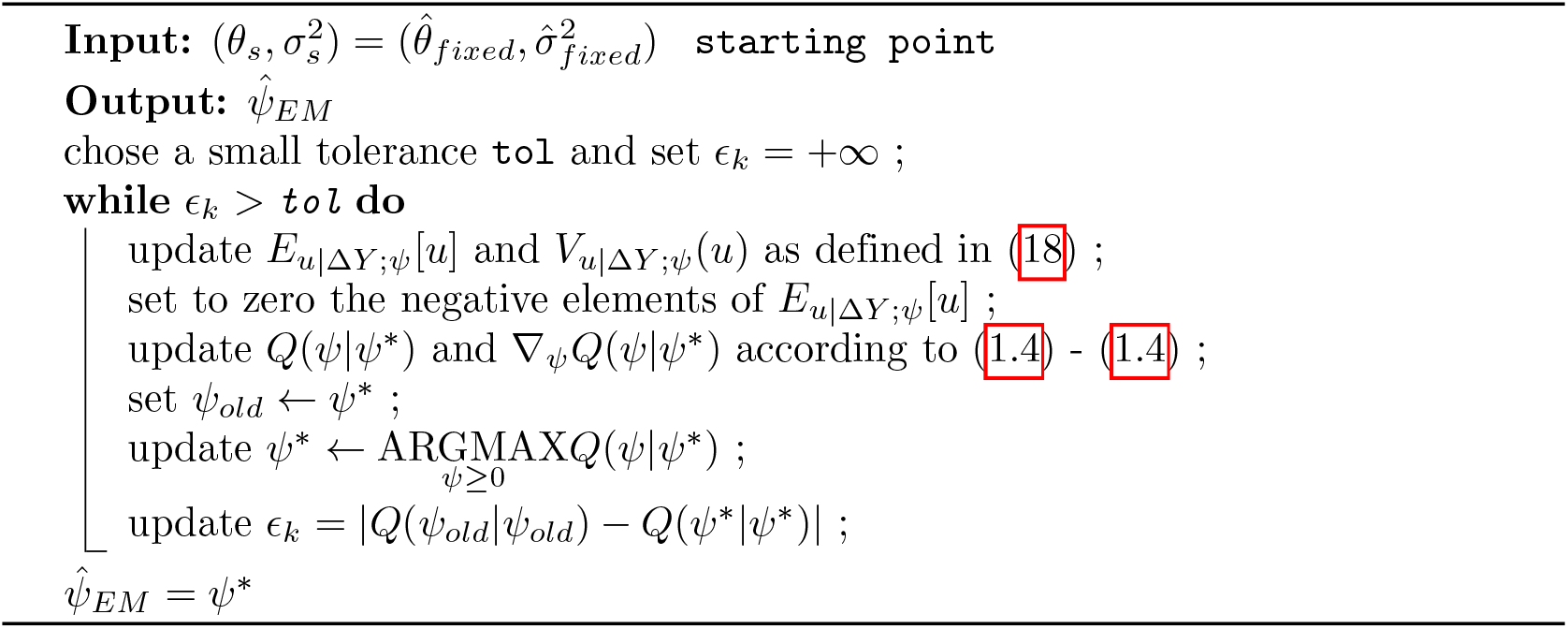

**Fig 1.**
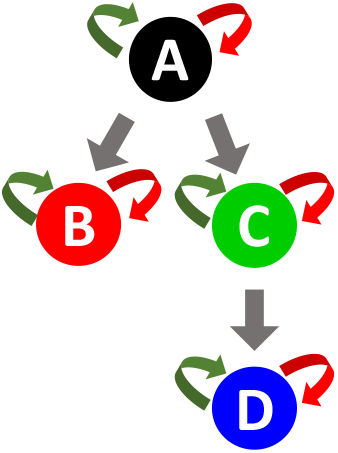
Differentiation structure of four synthetic cell types A, B, C, D. Duplication, death and differentiation moves are indicated with green, red and grey arrows respectively.

In the EM-algorithm we iteratively update the E-function *Q*(*ψ*|*ψ*\*) using the current estimate *ψ* * of *ψ* and then we minimize the − *Q*(*ψ*|*ψ*\*) w.r.t. *ψ* The EM algorithm is run until a convergence criterion is met, that is when the relative errors of both the E-step function *Q*(*ψ*|*ψ*\*) and the vector parameter *ψ* are lower than a predefined tolerance. Once we get the EM estimate 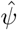 for the parameters we evaluate the goodness-of-fit of the mixed-model according to the conditional Akaike Information Criterion [3]. As every EM algorithm, the choice of the starting point *ψ*_s_ is very important from a computational point of view. We chose as a starting point 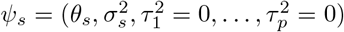 where 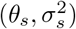 is the optimum found in the fixed-effects LLA formulation (24). This is a reasonable choice since we want to quantify how the dynamics 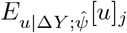 of each subject *j* departs from the average dynamics *θ*_*s*_. The EM pseudocode is given in Algorithm 3.

### 2 Simulation studies

Here we mimic the dynamics of *J* = 3 distinct clones in four synthetic cell types `A`, `B`, C, `D` following the differentiation network structure of Figure 4. The net-effect matrix *V* and hazard vector *h*(*Y, θ*) can be derived from equations (6)-(7) of the main paper. To simulate the clonal tracking data we used the *τ*-leaping Algorithm 1 with a time lag *τ* = 1, that has been run independently for each clone. We designed each simulation so that the first clone dominates lineage D and the third clone dominates lineage C. We first run a single simulation under different magnitudes for the noise variance *σ*^2^. Then we fitted both the base (24) and random-effects (27) models to the simulated data using Algorithms 2 and 3. We reported in Figure 2 the simulated trajectories and a scatterplot of the estimated conditional expectation 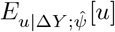 for the random-effects model against the true clone-specific parameters. In the same figure we also show a piechart where each clone *k* is identified with a pie whose slices are lineage-specific and weighted with *w*_*k*_, defined as the difference between the conditional expectations of the duplication and death parameters, that is 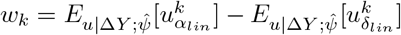 where `lin` is a cell lineage. The diameter of the *k*-th pie is proportional to the euclidean 2-norm of *w*_*k*_. Therefore, the larger the diameter, the more the corresponding clone is expanding into the lineage associated to the largest slice. The values of the estimated conditional expectations are reported in Table 1. The scatterplot of Figure 2 clearly indicate strong correlation between the true parameters and the conditional expectations 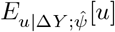 In particular, as expected, as the noise variance *σ*^2^ increases, the parameter estimates gradually move away from the diagonal, so that the correlation decreases. Also, our model correctly detected the dominance of clones 1 and 3 in lineages `D` and C respectively even for large values of *σ*^2^, as suggested by the pie-charts of Figure 2 and by the values of Table 1.

**Table 1.** Conditional expectations 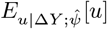 of the random-effects obtained from the estimated parameters 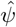 under different magnitudes of the noice variance *σ*^2^ (outer columns) for each clone (inner columns).

| | $\sigma^2 = 1$ | | | $\sigma^2 = 1$ | | | $\sigma^2 = 10$ | | | $\sigma^2 = 100$ | | |
| --- | --- | --- | --- | --- | --- | --- | --- | --- | --- | --- | --- | --- |
| | $c_1$ | $c_2$ | $c_3$ | $c_1$ | $c_2$ | $c_3$ | $c_1$ | $c_2$ | $c_3$ | $c_1$ | $c_2$ | $c_3$ |
| $\alpha_A$ | 0.183 | 0.191 | 0.198 | 0.183 | 0.191 | 0.198 | 0.151 | 0.139 | 0.127 | 0.000 | 0.105 | 0.050 |
| $\alpha_B$ | 0.146 | 0.148 | 0.145 | 0.146 | 0.148 | 0.145 | 0.163 | 0.148 | 0.137 | 0.442 | 0.337 | 0.358 |
| $\alpha_C$ | 0.163 | 0.168 | 0.518 | 0.163 | 0.168 | 0.518 | 0.166 | 0.175 | 0.649 | 0.300 | 0.214 | 0.958 |
| $\alpha_D$ | 0.450 | 0.100 | 0.121 | 0.450 | 0.100 | 0.121 | 0.479 | 0.199 | 0.319 | 0.499 | 0.691 | 0.778 |
| $\delta A$ | 0.001 | 0.001 | 0.001 | 0.001 | 0.001 | 0.001 | 0.001 | 0.000 | 0.001 | 0.000 | 0.000 | 0.000 |
| $\delta B$ | 0.007 | 0.007 | 0.007 | 0.007 | 0.007 | 0.007 | 0.008 | 0.007 | 0.007 | 0.016 | 0.014 | 0.015 |
| $\delta C$ | 0.004 | 0.004 | 0.004 | 0.004 | 0.004 | 0.004 | 0.004 | 0.005 | 0.005 | 0.008 | 0.006 | 0.008 |
| $\delta D$ | 0.002 | 0.002 | 0.002 | 0.002 | 0.002 | 0.002 | 0.002 | 0.003 | 0.004 | 0.002 | 0.010 | 0.007 |
| $\delta_{A \rightarrow B}$ | 0.129 | 0.130 | 0.133 | 0.129 | 0.130 | 0.133 | 0.127 | 0.126 | 0.110 | 0.119 | 0.177 | 0.173 |
| $\delta_{A \rightarrow C}$ | 0.148 | 0.149 | 0.151 | 0.148 | 0.149 | 0.151 | 0.154 | 0.155 | 0.153 | 0.197 | 0.197 | 0.133 |
| $\delta_{B \rightarrow C}$ | 0.079 | 0.080 | 0.078 | 0.079 | 0.080 | 0.078 | 0.082 | 0.079 | 0.058 | 0.081 | 0.084 | 0.059 |

Subsequently, to check parameter uncertainty we run *n*_*sim*_ = 100 independent simulations separately for each noise variance setting. After fitting both the base (24) and the random-effects (27) models, the latter always reached a significantly lower AIC compared to the null model as suggested by the boxplots of Figure 3. In Figure 3 we also report the boxplots of the estimated conditional expectation 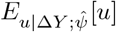, obtained from the independent simulations, divided by the true parameters *θ*_*true*_. Not surprisingly, as the noise variance *σ*^2^ increases, the parameter estimates get poor, but they still fluctuate around the true values, even under extreme magnitudes of *σ*^2^. These results clearly show accurate performance of the method in the identification of a simulated clonal dominance and in the inference of the true parameters, regardless of the noise level of the data.

**Fig 2.**
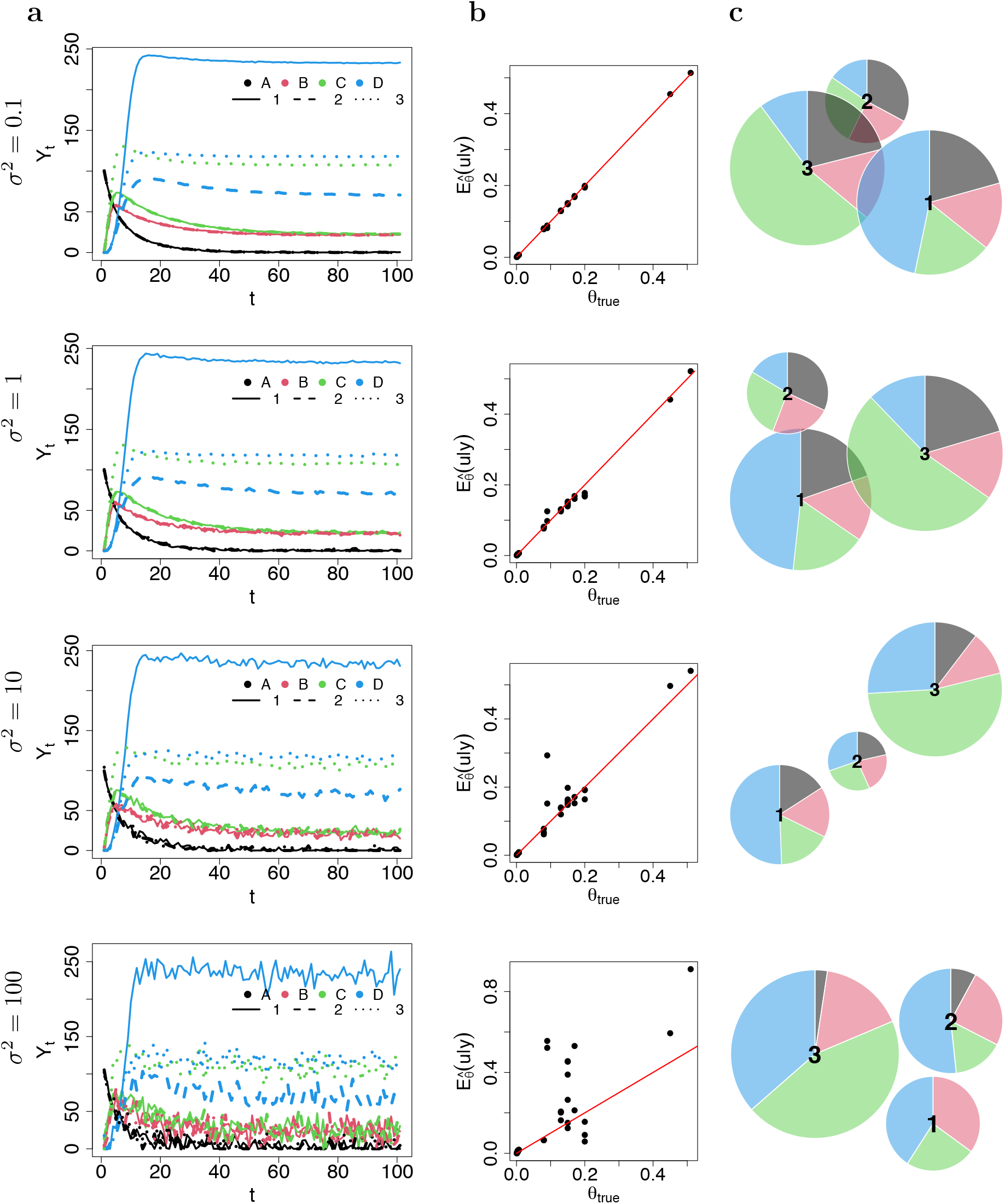
(a): Simulated trajectories. (b): Scatterplot between the clone-specific true parameters *θ*_*true*_ and the conditional expectation 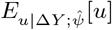 of the random effects obtained from the estimated parameters 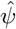 of the random-effects model. (c): Clonal pie-charts where each clone *k* is identified with a pie whose slices are lineage-specific and weighted with 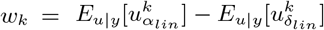. The diameter of the *k*-th pie is proportional to the euclidean 2-norm of *w*_*k*_. Each row refers to specific values of the noise variance *σ*^2^ used for simulations.

**Fig 3.**
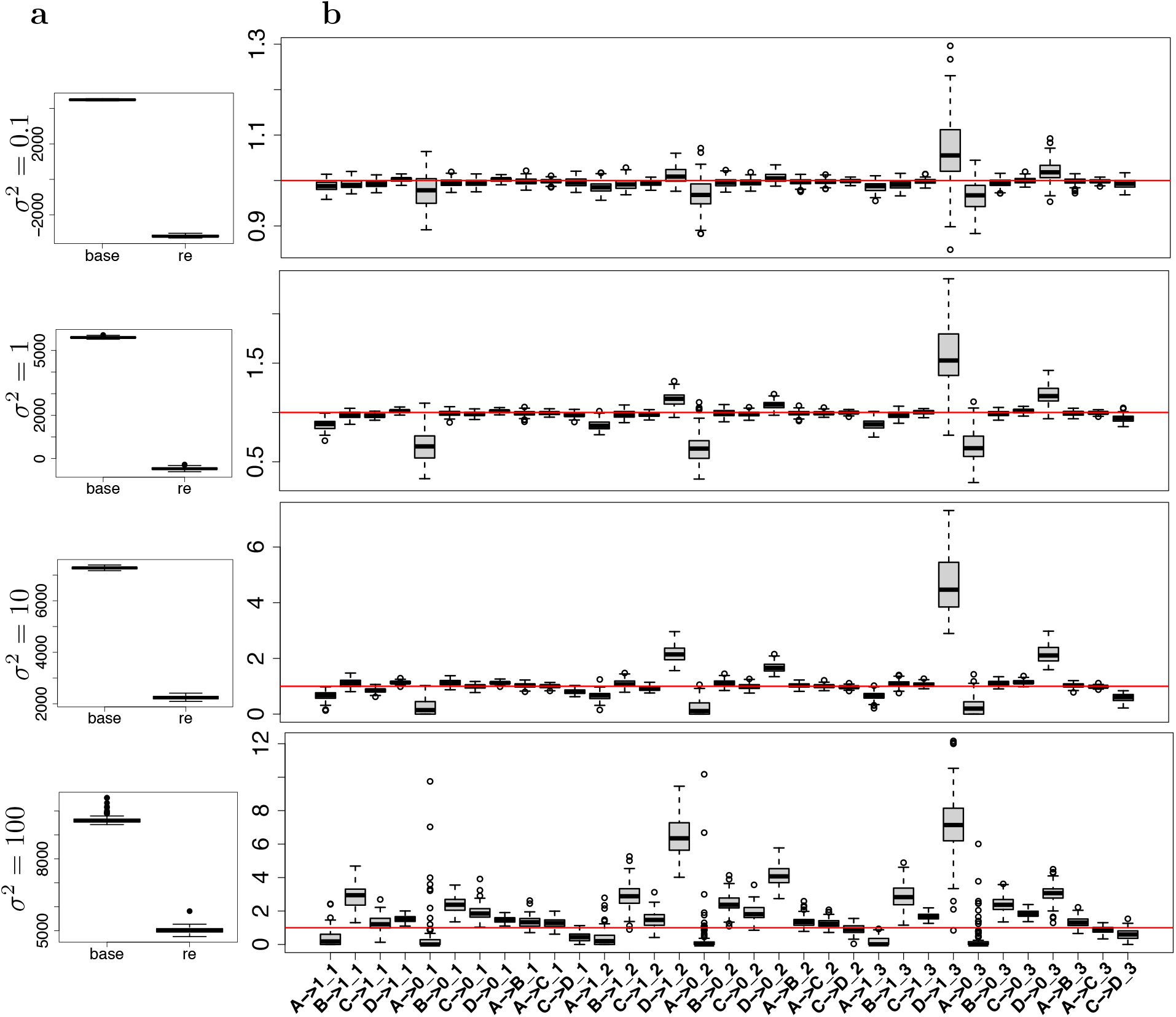
Boxplot of the AICs of the base and random-effect models (a) and boxplots of the estimated conditional expectation 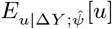 divided by the corresponding true parameters *θ*_*true*_ obtained under 100 independent simulations (b). Each row refers to a specific value of noise variance *σ*^2^ used for simulation.

### 3 Rhesus macaque data rescaling

Although the sample DNA amount was maintained constant during the whole experiment (200 ng for `ZH33` and `ZG66` or 500 ng for `ZH17`), the sample collected resulted in different magnitudes of total number of reads. Table 2 shows the total number of reads collected in each sample of the rhesus macaque clonal tracking dataset. This discrepancy makes all the samples not comparable across time and cell types. Therefore we rescaled the barcode counts according to

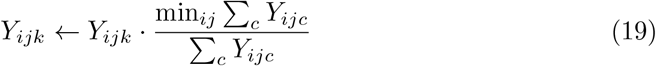

where *Y*_*ijk*_ is the *ijk*-entry of the barcode matrix with dimensions (*i, j, k*) mapping respectively time, cell type and clone.

**Table 2.** Total number of reads (sum across the different clones) collected in each treated animal at each time point and for all the cell types.

|  |  | T | B | NK | M | G |
| --- | --- | --- | --- | --- | --- | --- |
| ZH33 | 1 | 1465289 | 74735 | 135092 | 119331 | 2831 |
|  | 2 | 225797 | 216844 | 335789 | 1035270 | 908685 |
|  | 3 | 243986 | 413757 | 663184 | 886682 | 816990 |
|  | 4.5 | 485542 | 479493 | 834064 | 985821 | 987171 |
|  | 6.5 | 645005 | 676413 | 926089 | 895309 | 911637 |
|  | 9.5 | 829073 | 962325 | 1057398 | 1229233 | 1220506 |
| ZH17 | 1 | 51802 | 1347050 | 1288718 | 1351450 | 707382 |
|  | 2 | 826190 | 1342700 | 1350703 | 1354355 | 1213749 |
|  | 3 | 1303922 | 1347692 | 1338024 | 1347177 | 1283250 |
|  | 4.5 | 190591 | 1206361 | 489098 | 572877 | 1195585 |
|  | 6.5 | 887851 | 610999 | 1344488 | 381552 | 1339299 |
| ZG66 | 1 | 752127 | 0 | 211350 | 13382 | 0 |
|  | 2 | 692133 | 58890 | 308800 | 363310 | 145252 |
|  | 3 | 339292 | 209137 | 424458 | 808404 | 704331 |
|  | 4.5 | 617281 | 338977 | 718472 | 887183 | 897672 |

### 4 Estimated parameters

**Table 3.**
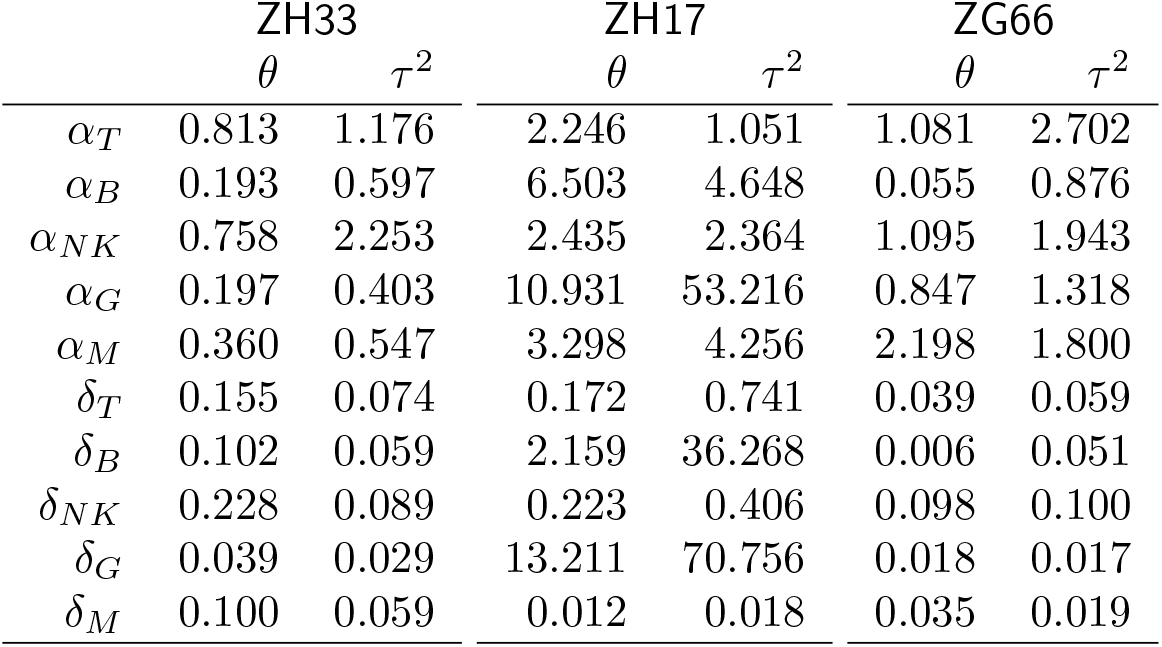
Parameter estimated for proposed mixed effects model: Fixed effects (*θ*) and variance (*τ* ^2^) of the random effects for both the duplication *α* and death *δ* parameters for each cell lineage and each treated animal.

**Fig 4.**
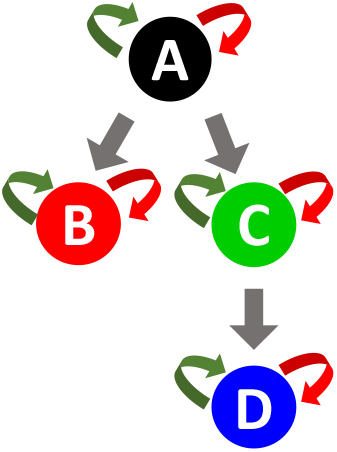
Cell differentiation structure of four synthetic cell types A, B, C and D. Duplication, death and differentiation moves are indicated with green, red and grey arrows respectively.

### 5 RestoreNet 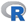 package: minimal working examples

This section reviews some key functionalities of `RestoreNet` package. Section 5.1 shows how to simulate a clonal tracking dataset from a stochastic quasi-reaction network. In particular, we show how to simulate clone-specific trajectories, following a given set of biochemical reactions. Sections 5.2 and 5.3 show how to fit the null (base) model and the random-effects model to a simulated clonal tracking dataset. Finally in Section 5.4 we show how to visualize the results at clonal level.

#### 5.1 Simulating clonal tracking datasets

A clonal tracking dataset compatible with `RestoreNet`’s functions must be formatted as a 3-dimensional array *Y* whose *ijk*-entry *Y*_*ijk*_ is the number of cells of clone *k* for cell type *j* collected at time *i*. The function `get.sim.tl()` can be used to simulate a trajectory of a single clone given an initial conditions *Y*_0_ for the cell counts, and obeying to a particular cell differentiation network defined by a list `rct.lst` of biochemical reactions. In particular, our package considers only three cellular events, such as cell duplication (*Y*_*it*_ → 1), cell death (*Y*_*it*_ → ∅) and cell differentiation (*Y*_*it*_ → *Y*_*jt*_) for a clone-specific time counting process

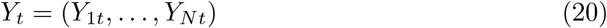

observed in *N* distinct cell lineages. The time counting process *Y*_*t*_ for a single clone in a time interval (*t, t* + Δ*t*) evolves according to a set of biochemical reactions defined as

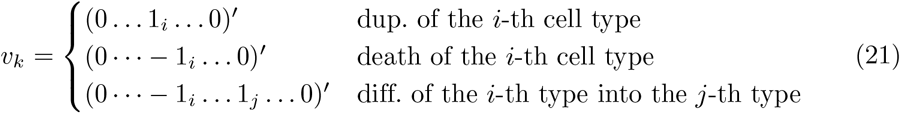

with the *k*-th hazard function given by

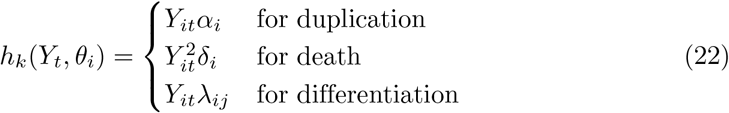

Finally, the net-effect matrix and hazard vector are defined as

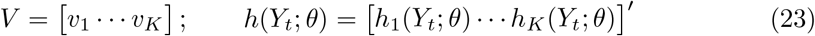

In particular, the cellular events of duplication, death and differentiation are respectively coded in the package with the character labels `“A->1”`, `“A->0”`, and `“A->B”`, where `A` and `B` are two distinct cell types. The following `R` code chunk shows how to simulate clone-specific trajectories of cells via a *τ*-leaping simulation algorithm. In particular, as an illustrative example we focus on a simple cell differentiation network structure of four synthetic cell types `A`, `B`, **C** and `D` and only one clone, as illustrated in Figure 4.

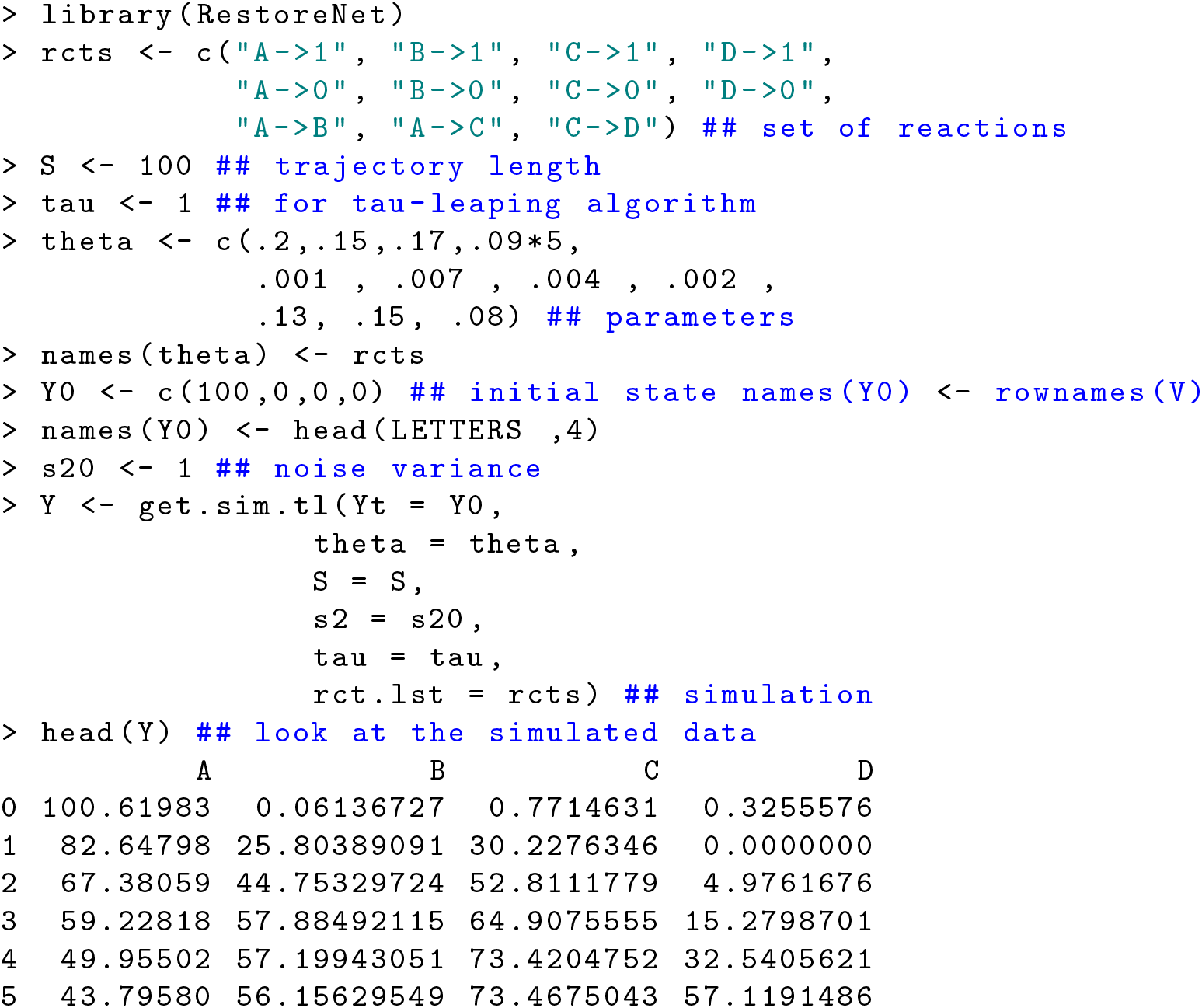

#### 5.2 Fitting the base model

The base model is defined as

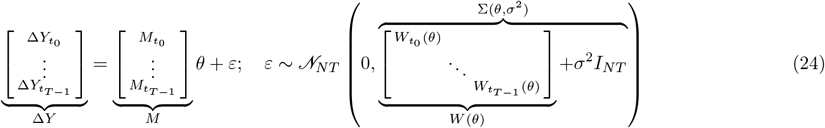

where

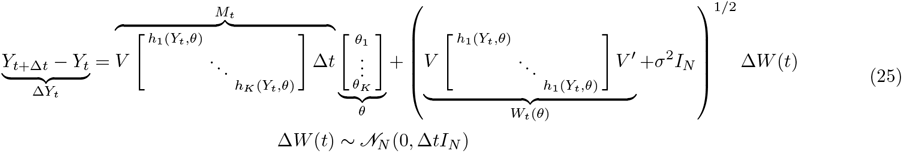

Further details can be found in []. The package `RestoreNet` allows to infer the parameters (*θ, σ*^2^) of (24) with a maximum likelihood approach, that is by solving the following constrained optimization problem

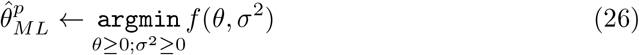

where the objective function *f* is the negative log-likelihood of the multivariate normal distribution *𝒩*_*NT*_ (*Mθ*, Σ (*θ, σ*^2^). The following `R` code chunk shows how to accomplish this on a clonal tracking dataset simulated from the same differentiation network structure of previous section. In this case we simulate the trajectories of three independent clones following different dynamics of clonal dominance, that is we use clone-specific values for the vector parameter *θ*.

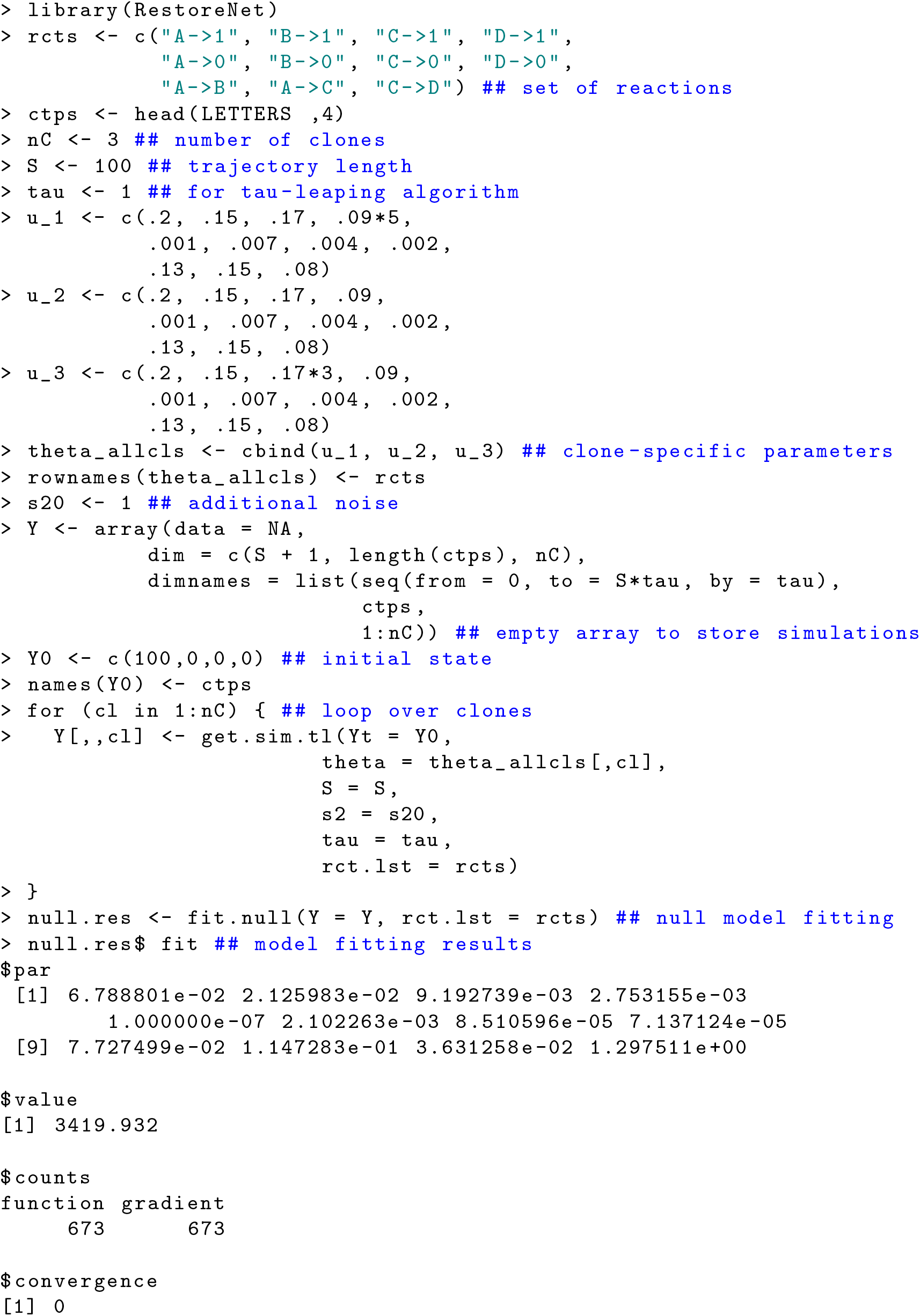
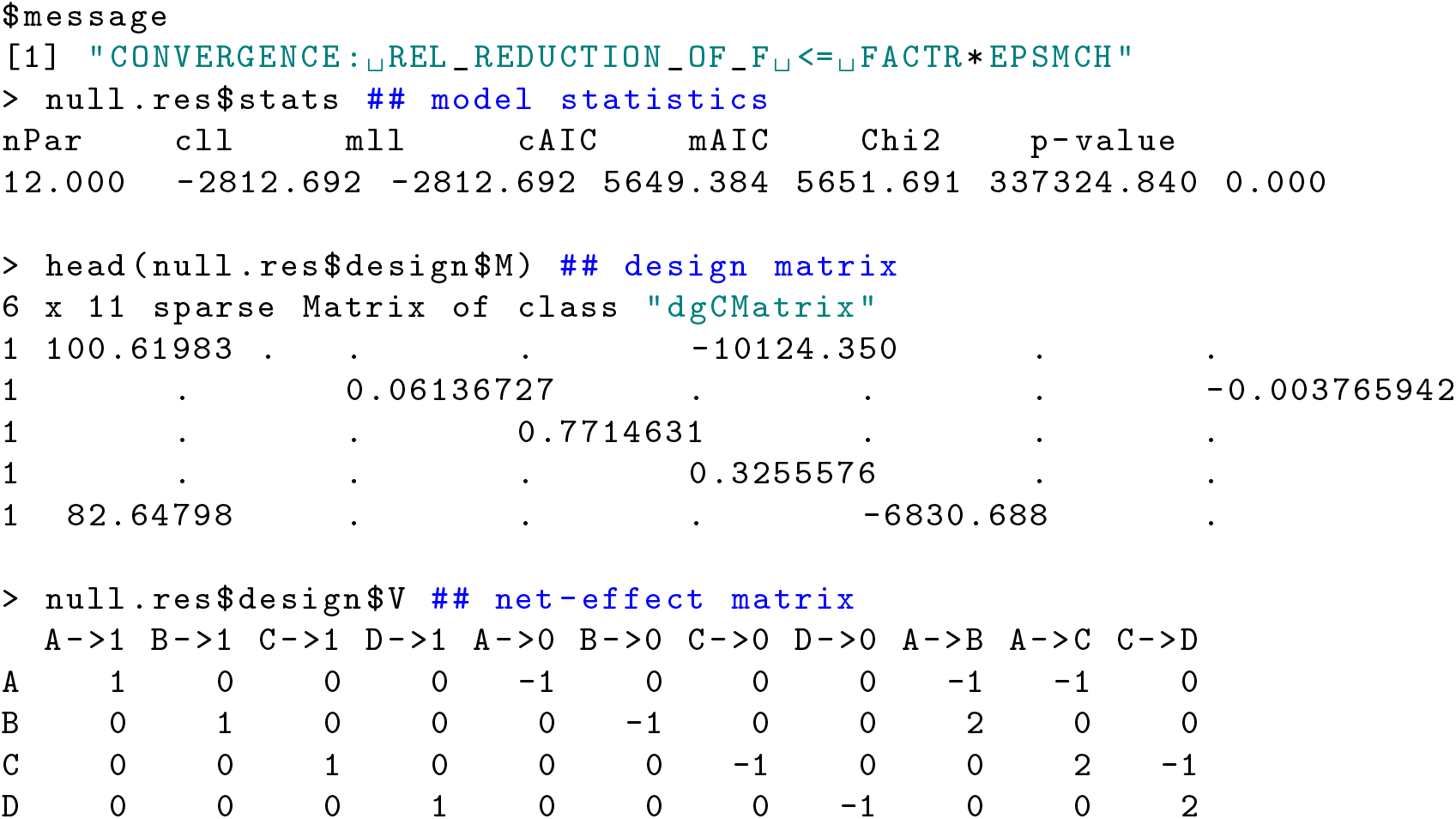

#### 5.3 Fitting the random-effects model

The random-effects model is defined as

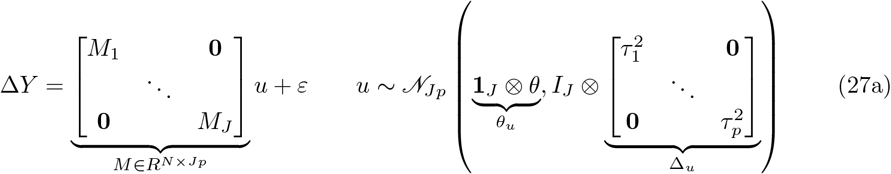

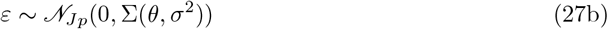

where *Y*_*t*+Δt_ − *Y*_*t*_ = Δ*Y* is the vector of cellular increments that took place in the time interval. Δ*t*, M is the block-diagonal design matrix for the random effects **u** centered in *θ, J* is the number of clones, and each block *M*_*j*_ is clone-specific. As in the case of the null model (24), to explain additional noise of the data, which has the additional advantage of avoiding singularity of the covariance matrix *W* (*θ*), we add to its diagonal a small quantity *σ*^2^ which we infer from the data. Under this framework (see [] for details) the conditional distribution of the random effects u given the data. Δ*Y* has the following explicit formulation

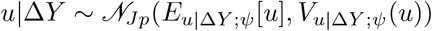

where *E*_*u*|Δ*Y* ; *ψ*_ [*u*] and *V*_*u*|Δ*Y* ;*ψ*_(*u*) provide clone-specific mean and variance of the (random) reaction rates. The package `RestoreNet` allows to infer the vector parameter 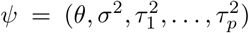, and in turn to get the corresponding conditional first two-order moments *E*_*u*|Δ*Y* ; *ψ*_ [*u*] and *V*_*u*|Δ*Y* ;*ψ*_(*u*) by the means of an efficient tailor-made Expectation-Maximization algorithm where Δ*Y* and *u* take the roles of the observed and latent states respectively. The following `R` code chunk shows how to accomplish this on the simulated clonal tracking dataset of previous section. In this example we use the optimal parameter vector 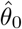 estimated for the null model in the previous section, as initial guess for the corresponding parameters in the random-effects model.

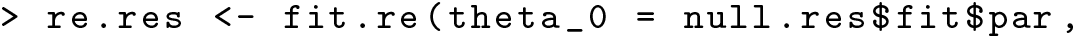
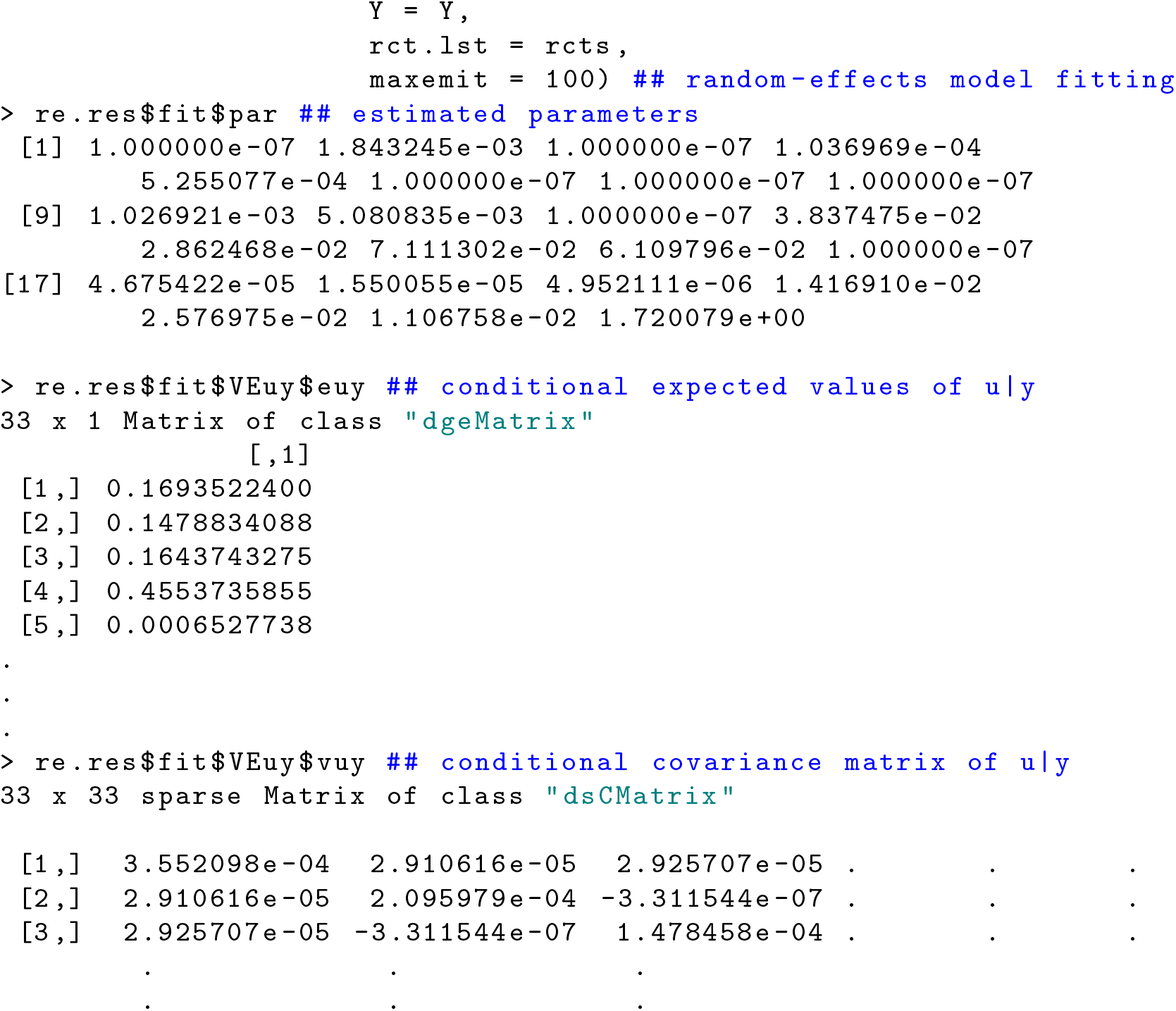

#### 5.4 Visualizing results

The main graphical output of `RestoreNet` is a clonal piechart. In this representation each clone *k* is identified with a pie whose slices are lineage-specific and weighted with *w*_*k*_, defined as the difference between the conditional expectations of the random-effects on duplication and death parameters, that is 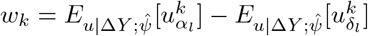, where *l* is a cell lineage. The diameter of the *k*-th pie is proportional to the euclidean 2-norm of *w*_*k*_. Therefore, the larger the diameter, the more the corresponding clone is expanding into the lineage associated to the largest slice. The package `RestoreNet` includes the function `get.scatterpie()` which returns a clonal piechart given a fitted random-effects model previously obtained with the function `fit.re()`. The following `R` code chunk illustrates how to obtain a clonal piechart with few lines of `R` code.

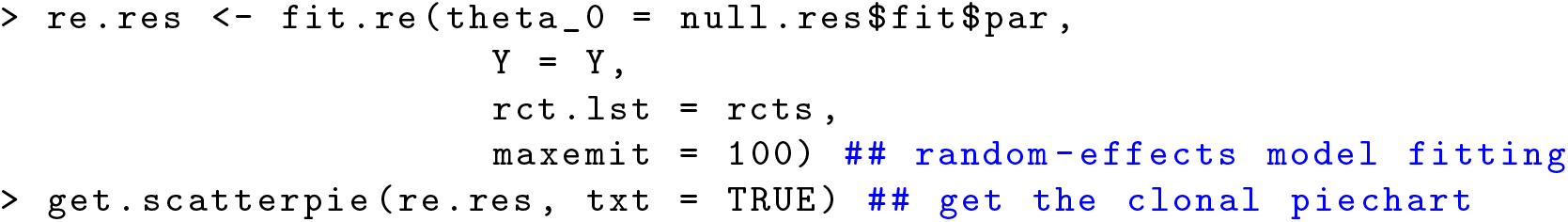

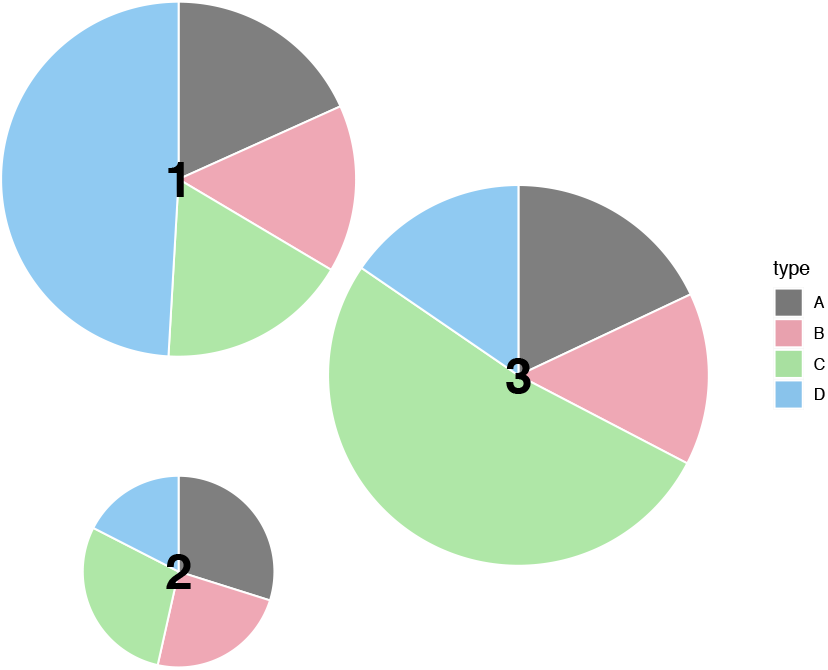

